# Metamorphic proteins at the basis of human autophagy initiation and lipid transfer

**DOI:** 10.1101/2022.09.21.508836

**Authors:** Anh Nguyen, Francesca Lugarini, Céline David, Pouya Hosnani, Barbora Knotkova, Anoshi Patel, Iwan Parfentev, Annabelle Friedrich, Henning Urlaub, Michael Meinecke, Björn Stork, Alex C. Faesen

**Affiliations:** Max-Planck Institute for Multidisciplinary Sciences, Laboratory of Biochemistry of Signal Dynamics, Göttingen, Germany; Institute of Molecular Medicine I, Medical Faculty and University Hospital Düsseldorf, Heinrich Heine University; Düsseldorf, Germany; Heidelberg University Biochemistry Center (BZH); Heidelberg, Germany; University Medical Centre Göttingen, Department of Cellular Biochemistry; Göttingen, Germany; Max-Planck Institute for Multidisciplinary Sciences, Bioanalytical Mass Spectrometry; Göttingen, Germany; University Medical Centre Göttingen, Institute of Clinical Chemistry, Bioanalytics Group; Göttingen, Germany

**Keywords:** autophagy, autophagy initiation, lipid transfer, membrane contact site, metabolism, protein metamorphosis

## Abstract

Autophagy is a conserved intracellular degradation pathway that uses de novo double-membrane vesicle (autophagosome) formation to target a wide range of cytoplasmic material for lysosomal degradation. In multicellular organisms, autophagy initiation requires the timely assembly of a contact site between the ER and the nascent autophagosome. Here, we report the in vitro reconstitution of a full-length sevensubunit human autophagy initiation supercomplex and found at its core ATG13-101 and transmembrane protein ATG9. Assembly of this core complex requires the rare ability of ATG13 and ATG101 to adopt topologically distinct folds. The slow spontaneous conversion between folds creates a rate-limiting step to regulate self-assembly of the super-complex. The interaction of the core complex with ATG2-WIPI4 enhances tethering of membrane vesicles and accelerates lipid transfer of ATG2 by both ATG9 and ATG13-101. Our work uncovers the molecular basis of the contact site and its assembly mechanisms imposed by the metamorphosis of ATG13-101 to regulate autophagosome biogenesis in space and time.

---

Macro-autophagy (called ‘autophagy’ hereafter) is an essential biological process of regulated degradation that promotes organismal health, longevity and helps combat cancer and neurodegenerative diseases (Mizushima et al., 2008). It is mediated by a specific cellular pathway that is capable of collecting cellular macromolecules as cargoes, including large aggregates and even organelles. The process is initiated by the nucleation of a flat membrane cisterna, termed ‘isolation membrane’ or ‘phagophore’, that then expands to surround the cargo and eventually closes to form the double-membrane autophagosome, which then fuses with the lysosome for the final degradation.

Recent work has shed light on the enigmatic molecular mechanisms that govern *de novo* autophagosome biogenesis. In multicellular organisms, the formation of autophagosomes requires the generation of contact sites between a subdomain of the ER (‘omegasome’) and the isolation membrane (Ktistakis, 2020). After the induction of autophagy, a complex array of proteins promptly colocalize at the contact site using unknown mechanisms (Fig. 1A). The early events of autophagy initiation involve three main protein components: the ULK complex (ULK1 or −2 kinase, FIP200 and HORMA domain proteins ATG13-101), the PI3-kinase complex (VPS34, VPS15, BECN1 and ATG14) and the trimeric transmembrane protein ATG9, a lipid scramblase that resides in highly dynamic small vesicles (Guardia et al., 2020; Maeda et al., 2020a; Mari et al., 2010; Orsi et al., 2012; Yamamoto et al., 2012; Young et al., 2006). These ATG9-containing vesicles are proposed to be seeds for membrane formation from the contact sites (Sawa-Makarska et al., 2020). Subsequently ATG2 and WIPI4 are recruited, which form the initial membrane tether to allow for lipids to flow into the growing autophagosome (Chowdhury et al., 2018; Maeda et al., 2019; Osawa et al., 2020; Osawa et al., 2019; Valverde et al., 2019). Interactions between subcomplexes, like ATG9 with ATG13-101 (Kannangara et al., 2021; Suzuki et al., 2015) and ATG2 (Gomez-Sanchez et al., 2018; Wang et al., 2001) or ATG13 with ATG14 (Jao et al., 2013; Park et al., 2016), have been found, but molecular details are lacking. It is unclear how these subcomplexes cooperate to form a functional contact site, how such a putative stable super-complex is assembled and disassembled, and how the different parts cooperate to perform the contact site function. To shed light on these questions and on additional functional and structural aspects of contact site function, we embarked on a reconstitution effort of a novel human autophagy initiation super-complex with purified full-length components.

**Figure 1:**
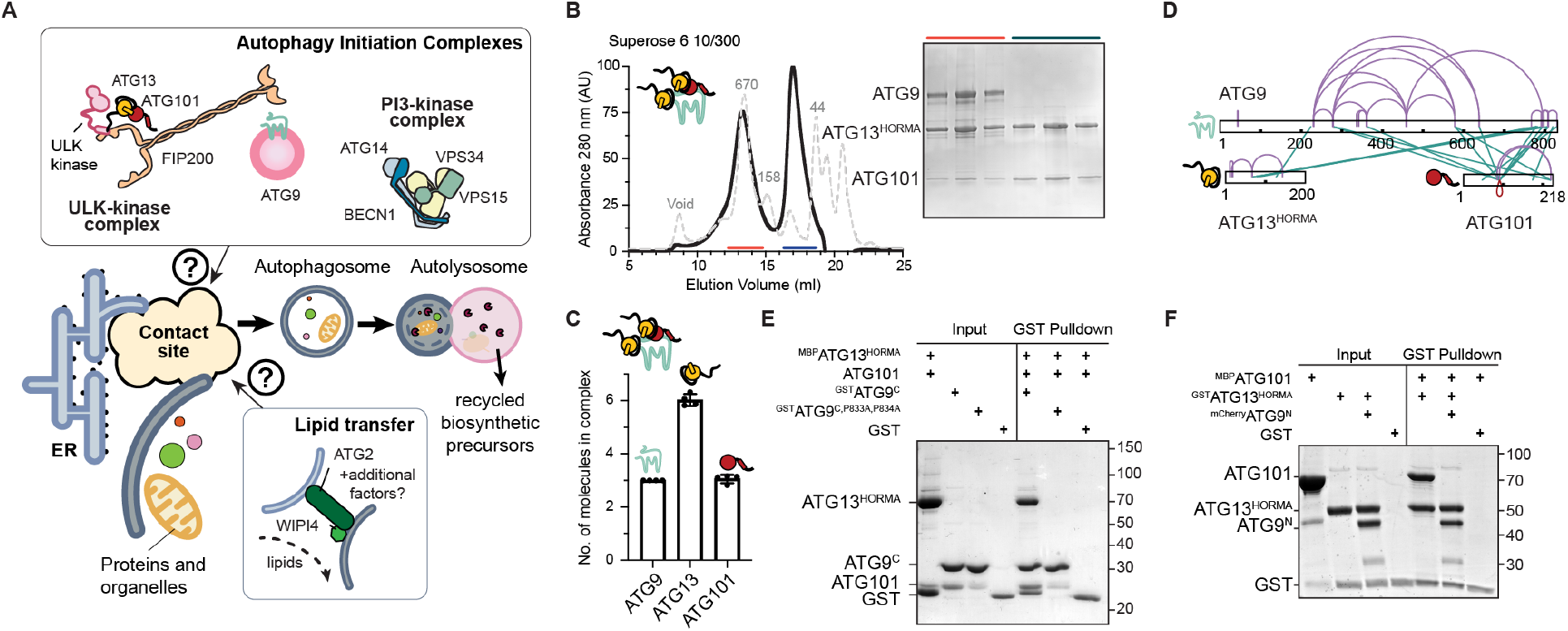
Reconstitution of ATG9-13-101 core complex. (**A**) The putative coalescence of early autophagy initiation complexes into a super-complex at the ER contact site promotes the *de novo* formation of an autophagosome. (**B**) ATG9, ATG13 and ATG101 form a stable complex. SEC profile of a complex of ^MBP^ATG9-^MBP^13^HORMA^-101 (red) and an excess of the ^MBP^ATG13^HORMA^-101 complex (blue) with corresponding fractions analyzed by SDS–PAGE. (**C**) Quantification of tryptophan fluorescence shows a 3:6:3 stoichiometry of the ATG9-13-101 complex. (**D**) Cross-linking coupled with mass spectrometry shows an extensive interaction interface between ATG9 and ATG13-101. (**E**) ATG13-101 interacts with ATG9^C^, which is lost after introducing P833A and P834A point mutants. ATG9^C^ (5 μM) and ATG13-101 (30 μM) were incubated at 4°C for 1 h before pull-down. (**F**) ATG101 does not interact with the ATG9^N^-ATG13^HORMA^ complex. Pull-down performed after 1 h incubation at 4 °C using 2 μM bait and 6 μM prey.

## Reconstitution of the ATG9-13-101 complex

The interaction between the ULK1 complex and ATG9 is one of the most upstream events in autophagy. To understand the molecular details of this interaction between ATG13-101 and ATG9, we purified the individual full-length proteins and the HORMA domain of ATG13 (ATG13^HORMA^) (Fig. S1A-D). As reported, ATG13 and ATG101 form a stoichiometric complex in solution (Qi et al., 2015) (Fig. S2A). Mixing ATG9 with an excess of the ATG13-101 complex, resulted in a stable ATG9-13-101 complex (Fig. 1B and S1E). Using the Stain-Free method, we determined a 3:6:3 stoichiometry of the ATG9-13-101 complex. This suggests a complicated assembly mechanism where the ATG13-101 dimer does not interact ‘en bloc’ (Fig.1C and Fig. S2A-C) (Gilda and Gomes, 2013). We observed that both ATG13 and ATG101 interact directly and stoichiometrically with ATG9 (Fig. S2D). Crosslinking experiments coupled with mass spectrometry, showed an extensive interface of both proteins with the C-terminal portion of ATG9 (Fig. 1D and Fig S1A). A structural model predicted by Alphafold2 (Jumper et al., 2021) shows that ATG9 residues 831 to 839 interact with both ATG13 and ATG101 (Fig S2E). Indeed, a construct that contained this region (ATG9^C^) interacts weakly with ATG13-101 and both proteins individually in stoichiometric fashion (Fig. 1E and Fig. S2F), while the ATG9^core^ transmembrane region could not (Fig. S2G). This interaction was lost when mutating the conserved residues P833 and P834 in ATG9^C^, confirming a recent independent structural analysis (Ren et al., 2022) (Fig. 1E and Fig. S2H). *Saccharomyces cerevisiae* Atg13 interacts with the poorly conserved N-terminal region of Atg9 (Suzuki *et al*., 2015). Although we observed no cross-links between ATG13 and ATG9^N^, ATG13 does interact stoichiometrically with ATG9^N^, while ATG101 does not (Fig. S2I,J). Overall, this reconstitution identified stoichiometric stable interactions between ATG9^N^ with ATG13, and ATG9C with the ATG13-101 dimer (Fig. S1A).

## Metamorphosis is rate-limiting for assembly

Surprisingly, ATG13 lost its ability to interact with ATG101 when bound to ATG9^N^ (Fig. 1F). Despite incubation with an excess of protein over 24 hours, ATG101 could not compete off ATG13, suggesting a non-canonical competitive interaction (Fig. 1F). ATG13 and ATG101 are HORMA (HOP1, REV7, MAD2) domain proteins known to use their rare metamorphic behavior to change their interaction spectrum (Gu et al., 2022). Metamorphosis allows them to switch between distinct conformer states under physiological conditions (Gu *et al*., 2022). The differential conformer interaction spectrum is pivotal for regulating the self-assembly of effector complexes. ATG13 and ATG101 are essential for autophagy and interact with multiple subcomplexes (Jao *et al*., 2013; Koyama-Honda et al., 2013; Wallot-Hieke et al., 2018), and are therefore prime candidates to control the assembly of a larger initiation machine.

Conformer switching is achieved by the unfolding and repositioning of structurally mobile elements (Fig. S1A). The emerging paradigm for HORMA domain proteins is that they default to an inactive state, before converting to a partner-bound active state. Client proteins are typically captured by wrapping the C-terminal ‘seatbelt’ around an interacting peptide, thereby creating a stable interaction. The considerable energy necessary for unfolding during metamorphosis, results in slow spontaneous conformer conversions (typically hours to days). Indeed, when mixing ATG13 or ATG101 with ATG9, we observed that complex formation was only achieved after an 18 to 24 hours incubation at 20 °C (Fig. 2A). So, both ATG13 and ATG101 default to an inactive non-ATG9-binding state, and that a conformer switch is obligatory before interacting with ATG9. In contrast, ATG13 interacts with ATG101 instantaneously, so the default conformer of ATG13 requires no metamorphosis to interact with ATG101 (Fig. S3A).

**Figure 2:**
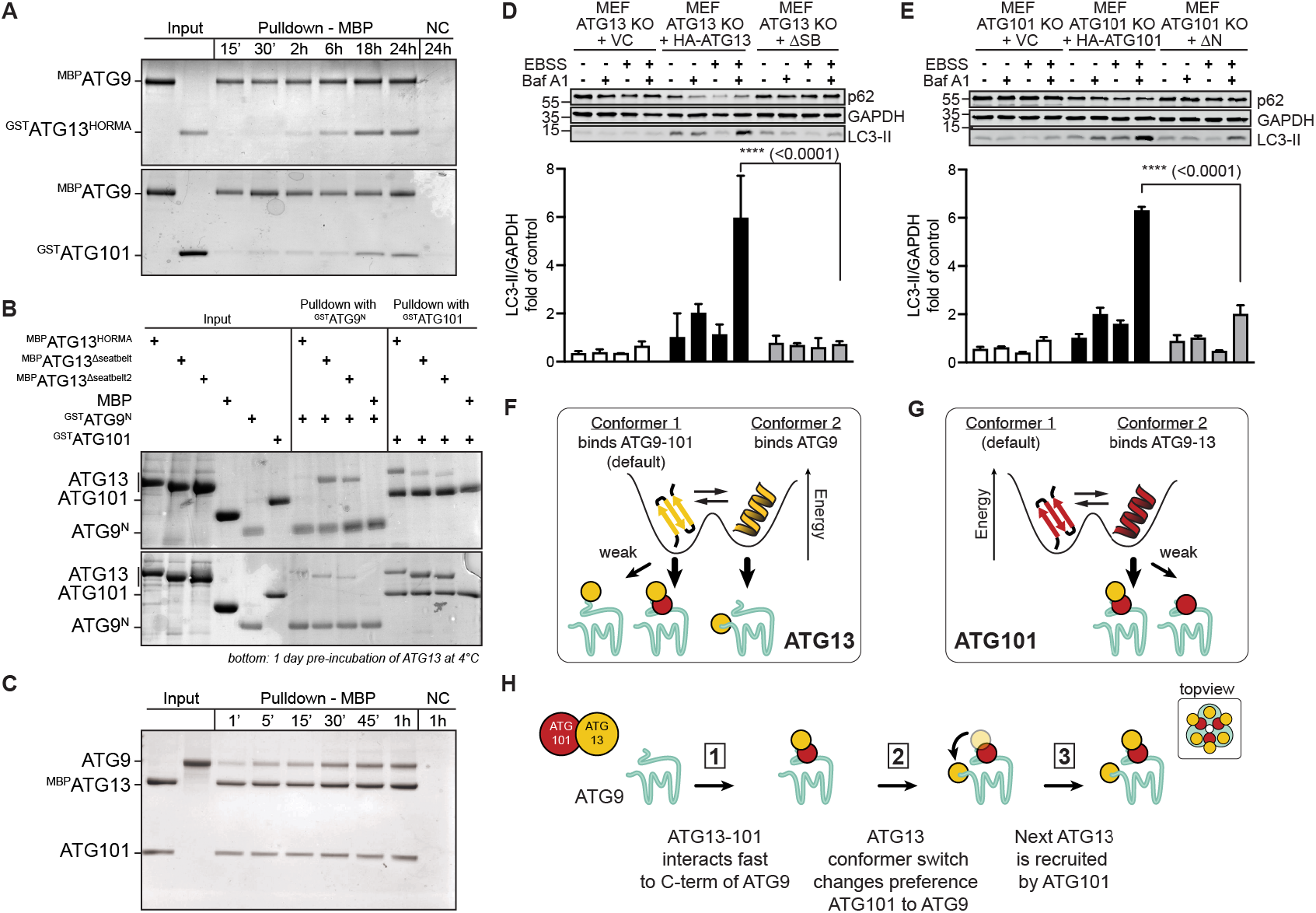
ATG13 and ATG101 metamorphosis dictates assembly of the core complex. (**A**) Assembly of ATG9-13 (top) and ATG9-101 (below) takes 18-24 hours. Pull-down performed after incubation at 20°C using 2 μM protein. (**B**) ATG13 conformers bind ATG9^N^ or ATG101. ATG13^WT^ quickly interacts with ATG101, while mutants default to the ATG9^N^-binding conformer. After incubation at 4°C without binding partners, the ATG13 mutants prefer binding to ATG101. Pull-down performed after 1 h incubation using 1 μM bait and 3 μM prey. (**C**) Assembly of ATG9-13-101 takes 30 to 60 minutes. Pull-down performed after 1 h incubation at 4 °C using 0.1 μM bait and 0.2 μM prey. (**D**,**E**) ATG13 and ATG101 conformer mutants decrease basal and starvation-induced LC3 and p62 turnover in MEFs. KO MEFs retrovirally transfected with empty vector (VC) or ATG13 or ATG101 cDNA variants were incubated for 2 h in growth or starvation medium (EBSS) with or without 40 nM Bafilomycin A_1_. ΔSB =△seatbelt (Fig. S1A). Results are mean + SD. Statistical analysis used two-way ANOVA (using Tukey’s multiple comparisons test). (**F**,**G**) Both ATG13 (F) and ATG101 (G) can adopt two conformers with each an exclusive interaction spectrum. (**H**) Hand-over assembly model of the ATG9-13-101 complex.

Removing mobile elements in HORMA domains can change the default conformer state (Mapelli and Musacchio, 2007). We therefore created ATG13 mutants that lack either the N-terminal (ATG13^ΔN^) or the C-terminal (ATG13^Δseatbelt^) mobile elements, or that contain a shortened internal loop predicted to allow for switching (ATG13^LL^, for ‘loop-less’) (Fig. S1A). All these ATG13 mutants lost the ability to bind to ATG101 within one hour (Fig. S3B). However, in presence of ATG101, ATG13^Δseatbelt^ will slowly convert to an ATG101-binding competent conformer (Fig. S3A). To confirm that ATG13 can indeed adopt two states with differential binding capabilities, we tested the mutants preferential binding to either ATG9^N^ or ATG101 after relatively short incubations (1 hour). We observed that ATG13^WT^ prefers to bind to ATG101, while the ATG13 mutants preferred ATG9^N^ (Fig. 2B, top). Following a path previously explored with other metamorphic proteins, we aimed to induce a conformer switch by incubating the ATG13 constructs overnight at reduced temperatures (4 °C) in the absence of binding partners. This did not affect the preference of ATG13^WT^, which still bound readily to ATG101 but not ATG9^N^ (Fig. 2B, bottom). In contrast, the mutants did switch preference from ATG9^N^ to ATG101. Overall, this shows that the thermodynamically most stable ATG13^WT^ conformer prefers to bind ATG101 and that a slow conformer change is required to bind to ATG9^N^.

Metamorphic proteins can induce conformational switching through dimerization, which can be crucial in accelerating interaction kinetics of HORMA domains with their client (Gu *et al*., 2022). Since dimerization of ATG13 and ATG101 is essential for autophagy (Wallot-Hieke *et al*., 2018), we wondered if dimerization of ATG13 and ATG101 affects their interaction kinetics to ATG9. Indeed, we observed that their complex formation with ATG9^FL^ or ATG9^core+C^ is accelerated to 30 minutes (Fig. 2C and Fig. S3C). In contrast, the interaction of ATG13 to ATG9^N+core^ is not accelerated by the presence of ATG101 (Fig. S3C). This shows that dimerization of ATG13 and ATG101 dramatically accelerates their interaction to ATG9^C^, while the interaction of ATG13 to ATG9^N^ remains remarkably slow.

Next, we wondered what consequence the conformer de-stabilizing mutants would have in cells. To probe autophagic degradation activity, we measured autophagic flux in murine embryonic fibroblasts (MEFs) (Klionsky et al., 2021). Upon inducing starvation by treating the cells with EBSS, both LC3 and p62/SQSTM1 turnover was rescued in cells expressing the wild-type proteins in ATG13 or ATG101 knock-out cells (Fig. 2D,E and Fig. S3D). However, autophagic flux was largely abolished in cells expressing either ATG13^Δseatbelt^ or ATG101^ΔN12^. Since the ATG13 mutants affect the conformer states and therefore the interaction with ATG101, we also investigated the stability of the ULK1 complex. ATG101 was not stabilized in cells expressing ATG13^Δseatbelt^, nor were ULK1 and ATG13 stabilized by ATG101^ΔN12^, confirming that these mutants likely abolish the dimerization between ATG13 and ATG101 resulting in the destabilization of the ULK complex (Fig. S3E). In cells expressing ATG13^Δseatbelt^, ULK1 was stabilized as compared to the parental *Atg13* knockout MEFs. However, the apparent molecular weight of ULK1 was lower than in cells expressing wild-type ATG13, indicating altered post-translational modifications (Fig. S3E). In cells expressing ATG13^Δseatbelt^, inhibitory phosphorylation at ULK1 Ser757 and Ser637 was reduced compared to wild-type ATG13-expressing cells, but remained responsive towards starvation (Fig. S4A). This suggests that the autophagy initiation pathway upstream of ULK1 is functional (Wong et al., 2015). In contrast, starvation-induced and ULK1-dependent phosphorylation of ATG14 at Ser29 was completely abolished in these cells, indicating a defective autophagy signaling downstream of ULK1 (Wold et al., 2016). Notably, in cells expressing the ATG101^ΔN12^ variant, both ULK1 activation and activity appeared unaffected compared to wildtype ATG101 (Fig. S4B), although this truncation clearly affected autophagic flux. Collectively, these observations suggest that changing the default conformer states of either ATG13 and ATG101 abolishes autophagic flux. Autophagic signaling downstream of ULK1is most sensitive to mutating ATG13, which strongly reduces binding to ATG101 leading to its instability.

Overall, these results can be summarized in the following ‘hand-over model’ (Fig. 2F-H). ATG13 and ATG101 can interact with ATG9, but this requires a slow obligatory conformer switch (Fig. 2F,G). ATG13 defaults to a state that binds ATG101 (Fig. 2F), and together they create the composite interface that allows for quick interaction with ATG9^C^ (Fig. 2H, step 1). Here, ATG13 can switch conformer state, and ATG101 ‘hands over’ the ATG13 molecule to ATG9^N^ (Fig. 2F and H, step 2). The increased local concentration and proximity might aid in this process. Next, ATG101 can recruit another ATG13 to saturate all ATG9 molecules in the trimer (Fig. 2H, step 3). This would yield a 3:6:3 stoichiometric complex, although sub-stoichiometric amounts of ATG101 might suffice to saturate all ATG9 monomers with ATG13.

## ATG9-13-101 as an interaction hub for supercomplexes

Next, we asked if an assembled ATG9-13-101 complex could bring all the early autophagy initiation complexes together in a stable supercomplex. To test this, we purified recombinant fulllength human ULK1, FIP200, ATG14 and BECN1 (Fig. 3A and Fig. S5A-C). ATG13-101 could form a stable stoichiometric complex when mixed with full-length ULK1 and FIP200 to form the ULK1 complex (Fig. 3B and Fig. S5D). ATG13-101 also interacted with ATG14-BECN1 (Fig. 3B and Fig. S5E). When we immobilized ULK1, we could specifically precipitate a stable 7-subunit supercomplex containing the ULK1 complex, the ATG9-13-101 complex and ATG14-BECN1 from the PI3-kinase complex (Fig. 3C). When omitting ATG9-13-101, the super-complex did not form, highlighting the coordinating role of the core complex in the assembly (Fig. S5F).

**Figure 3:**
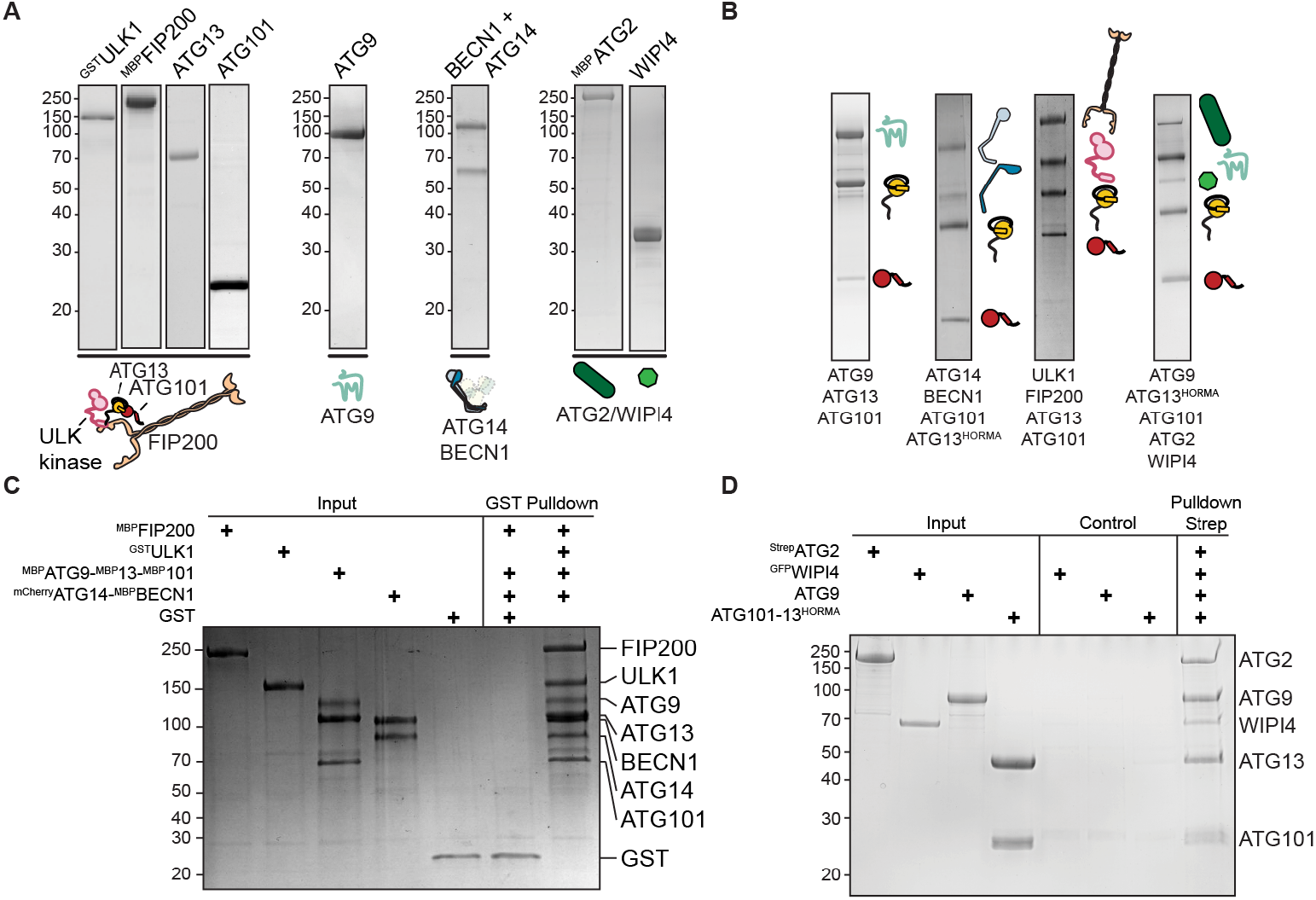
The ATG9-13-101 complex is a core interaction hub. (**A**,**B**) Coomassie-stained SDS–PAGE gallery of recombinant individual proteins (A) and ATG13-101 containing complexes (B) used in this study. The complexes constitute the core ATG9-13-101 complex, the ULK1 complex, the PI3-kinase subunits ATG14-BECN1 interacting with the HORMA domains of ATG13-101 and the complex between the ATG9-13-101 core complex and lipid transfer complex ATG2-WIPI4. (**C**) *In vitro* GST-ULK1 pull-down experiment showing a stable and defined 7-subunit complex containing almost all the full-length proteins of the canonical autophagy initiation complexes: ATG9-13-101-FIP200-ULK1-ATG14-BECN1. Pull-down was performed using 1 μM bait and 3 μM prey, with the proteins were incubated at room temperature for 1 h. (**D**) *In vitro* pull-down experiment using ^strep^ATG2 as bait showing the stable incorporation of ATG2-WIPI4 and ATG9-13-101 in a 5-subunit complex. Pull-down was performed using 1 μM bait and 3 μM prey, with the proteins were incubated at room temperature for 1 h.

ATG9 interacts with ATG2 which is proposed to establish membrane contact sites, together with the ATG2-adaptor protein WIPI4/Atg18 (Ghanbarpour et al., 2021; Gomez-Sanchez *et al*., 2018; Tang et al., 2019; Wang *et al*., 2001). We therefore wondered if the ATG9-13-101 complex could also accommodate ATG2 and WIPI4. We purified ATG2 and WIPI4, which form a stable complex (Fig. 3A and Fig. S5G-J) (Lu et al., 2011). While WIPI4 showed no interaction with ATG9 (Fig. S5J,K), ATG2 could directly interact with both to form a defined complex in solution (Ext. data Fig. 5I; black) as well as in pulldown experiments (Fig. S5K). Next, we used ATG2 as a bait to combine all proteins into a 5-subunit ATG2-WIPI4-ATG9-13-101 complex (Fig. 3D and Fig. S5L). We also note the increased intensity of ATG9-13-101 compared to ATG2-WIPI4, suggesting that an ATG2 monomer likely interacts with an ATG9 trimer (Fig. 3B,D). Overall, this shows that the ATG9-13-101 complex can combine both the ULK1 complex and PI3-kinase complex in a stable super-complex, as well as be part of a larger lipid transfer complex with ATG2.

**Figure 4:**
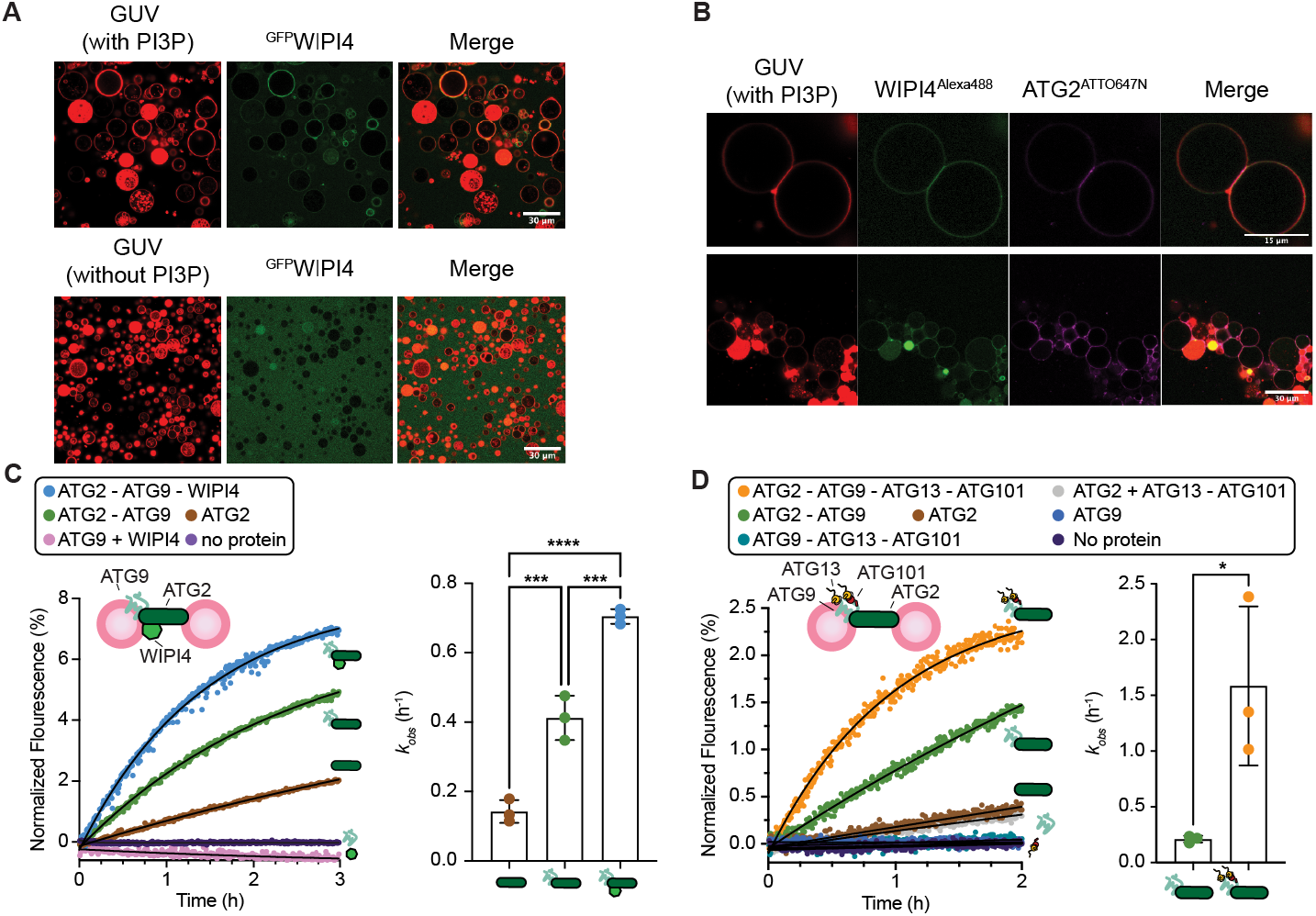
Lipid transfer is accelerated by both ATG9 and ATG13-101. (**A**,**B**) Representative images show the membrane localization of lipid transfer complex ATG2-WIPI4. ^GFP^WIPI4 (200 nM; green) bound to Rh-GUVs (red) in a PI3P-dependent manner (A). WIPI4 (200 nM; green) and ATG2 (200 nM; purple) cooperate to tether GUVs (red) to form an extensive membrane contact site (B). Representative images show the tethering of two (top panel) or a cluster of GUVs (bottom panel). (**C**) FRET-based lipid transfer assay showing the effects of ATG9 (100 nM) and WIPI4 (100 nM) on the lipid transfer efficiency of ATG2 (33 nM). Statistical significance was determined by one-way ANOVA followed by Turkey’s multiple comparisons test. All values are mean ± SD; ***, P < 0.001; ****, P < 0.0001. The experiments were performed at 25 °C with 3 independent technical replicates. A representative dataset is shown. (**D**) FRET-based lipid transfer assay showing the effects of ATG13^HORMA^-ATG101 (100 nM) and ATG9 (100 nM) on lipid transfer efficiency of ATG2 (33 nM). Statistical significance was determined by Student’s t-test. All values are mean ± SD; *, P ≤ 0.05. The experiments were performed at 25°C with 3 independent technical replicates. A representative dataset is shown.

**Figure 5:**
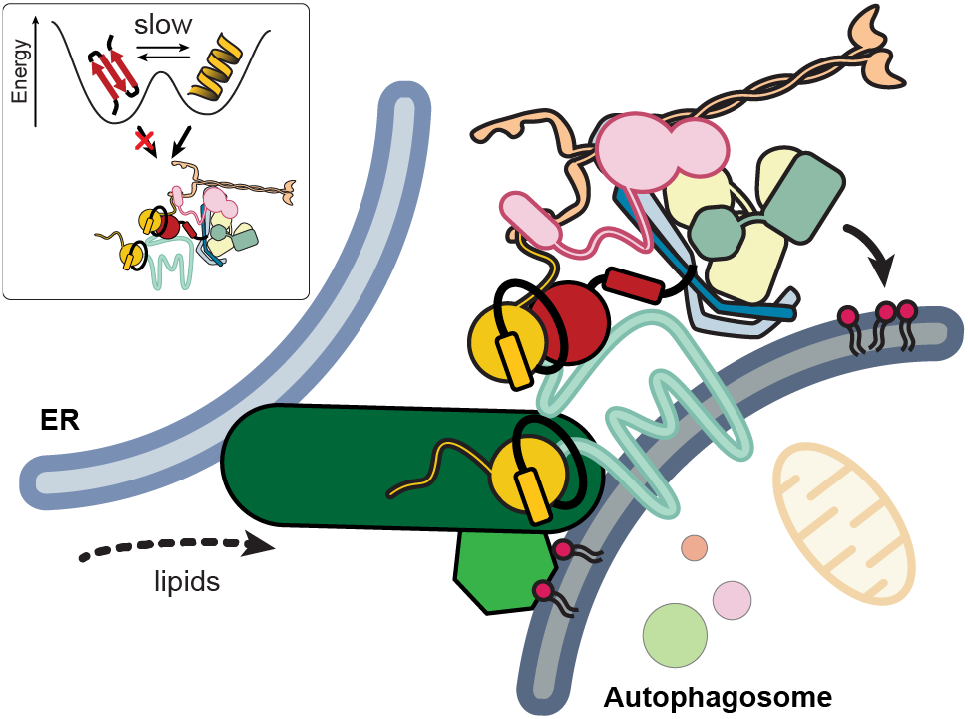
Summary of our findings. All functional subcomplexes form a defined and stable super-complex. The ATG9-13-101 complex is an interaction hub, whose assembly represents a rate-limiting intermediate due to the slow obligatory conversion between topologically distinct native folds of ATG13 and ATG101 (inset).

## Acceleration of ATG2-dependent lipid transfer

Next, we wondered if this larger assembly would affect the membrane tethering and lipid transfer activity of ATG2. WIPI4 is an ATG2-adaptor protein that binds the phospholipid phosphatidylinositol 3-phosphate (PI3P or PtdIns3P), which is part of the lipid signature of autophagic membranes (Burman and Ktistakis, 2010). To visualize membrane binding and tethering by the ATG2-WIPI4 complex using confocal microscopy, we created fluorescently labeled ATG2 and WIPI4. WIPI4 binds strongly and specifically to GUVs containing PI3P, but we observed no tethering (Fig. 4A). Upon addition of ATG2 to WIPI4, we observed of ATG2 to membranes resulting in a striking clustering of vesicles (Fig. 4B, bottom panels) and the establishment of extensive contact between vesicles that show enriched levels of ATG2-WIPI4 (Fig. 4B, top panels).

Next, we used those conditions to test the ATG2-tethering capabilities of large unilamellar vesicles (LUVs) using dynamic light scattering (DLS). The apparent particle size increased upon adding ATG2, but not when adding ATG9 or WIPI4 (Fig. S6D,E). Adding the non-specific proteinase K reverted this effect, showing that ATG2 can reversibly tether vesicles (Fig. S6D,E). Adding WIPI4 to ATG2 increased the particle size further, likely by cooperatively improving the membrane binding of ATG2. Next, we successfully reconstituted ATG9 into protein-free LUVs (PLs) as judged by the floatation of ATG9 to the top fractions of a Nycodenz gradient (Fig. S6C). Using a protease protection assay, we observed that the N-terminal MBP tag could be removed from ATG9, indicating that almost all ATG9 was properly oriented in the vesicle bilayer (Fig. S6D). Using the ATG9 PLs together with ATG2-WIPI4, we observed a further increase in particle size using DLS, which could be reverted to sizes similar to non-tethered LUVs after incubation with proteinase K (Fig. S6E). Overall, this shows that the ATG2 tethering capabilities are cooperatively enhanced by both ATG9 and WIPI4.

As a peripheral membrane protein, ATG2 mediates lipid transfer between the outer cytosolic membrane leaflets of donor and acceptor vesicles. This results in an asymmetric lipid distribution between the outer and the inner luminal leaflets. The scramblase activity of ATG9 might aid the lipid transfer of ATG2 by re-equilibrating lipids between the leaflets and avoid membrane destabilization (Gomez-Sanchez *et al*., 2021; Maeda *et al*., 2020b; Noda, 2021; Reinisch *et al*., 2021). We confirmed that reconstituted ATG9 in PLs shows robust scramblase activity as judged by the increased bleaching of fluorescently labelled lipids by dithionite compared to ATG9-free liposomes (Maeda *et al*., 2020b; Matoba et al., 2020) (Fig. S6E). In cells, ATG9 is suggested to reside in the acceptor membrane, while other scramblases reside on the donor membrane on the ER (Ghanbarpour *et al*., 2021). To test the effect of adding scramblases to membranes, we measured lipid transfer by ATG2 between two ATG9 containing PLs (Fig. S6F). In this established assay, donor LUVs are prepared with fluorescent lipids at a concentration sufficient for FRET (2% NDB-PE and 2% Rhodamine-PE) (Maeda *et al*., 2019; Osawa *et al*., 2020; Osawa *et al*., 2019; Valverde *et al*., 2019). After adding a non-fluorescent acceptor LUV, lipid transfer is quantified by the increase of fluorescence of NDB due to the reduction in FRET after lipids flow between LUVs. The addition of ATG9 increased lipid transfer rate of ATG2 by approximately 3 times (with observed rates of 0.14 h^-1^ and 0.4 h^-1^, respectively), while adding both ATG9 and WIPI4 increased the lipid transfer of ATG2 approximately 5 times (k_obs_ of 0.7 h^-1^) (Fig. 4C).

The ATG2-WIPI4 and ATG9-13-101 complexes stably interact, but the molecular details of this complex are unclear (Fig. 3B,D and Fig S5L). The apparent 1:3 stoichiometry of ATG2 to ATG9 suggests that ATG2 interacts in close proximity to the ATG9 lipid entry and exit pores. ATG13 and ATG101 crosslinked to the core of ATG9, which could feasibly modulate lipid transfer (Fig. 1D). We pre-assembled the ATG9-13-101 complex by incubating ATG13^HORMA^-101 with ATG9 proteo-liposomes, before adding ATG2 to start the lipid transfer. Compared to ATG2-ATG9, the presence of ATG13-101 increased the lipid transfer rate approximately 7.8 times (k_obs_ of 1.57 h^-1^ and 0.2 h^-1^, respectively) (Fig. 4D). Overall, this shows that both ATG9 and ATG13-101 cooperatively enhance the lipid transfer rate of ATG2 by over 20-fold in these conditions.

## Discussion

After the induction of macro-autophagy, a membrane contact site is assembled between the ER and the *de novo*-formed autophagosome that colocalizes the components of the autophagy initiation machinery. Here, we have presented the production of two novel super-complexes: a 7-subunit super-complex that stably comprises almost all of the fulllength proteins of the early human autophagy initiation components (the ULK1 complex, ATG9 and ATG14-BECN1 of the PI3-kinase complex) and a 5-subunit super complex with ATG2/WIPI4 (Fig. 5). They share at their core the ATG9-13-101 complex, which serves three functions: 1) the spontaneously but slow metamorphosis of HORMA domain proteins ATG13 and ATG101 creates an obligatory rate-limiting step in its assembly, 2) it coordinates the co-incidence of all subcomplexes in a stable super-complex, and 3) it interacts with ATG2-WIPI4 and accelerates its lipid transfer rate. This is the first time that HORMA domains have been shown to not only control the assembly of an effector complex, but also to modulate its activity.

The molecular basis for a regulated assembly of the initial contact site between the ER and the newly formed autophagosome is a fundamental unresolved issue. We show that the super-complexes are thermodynamically stable, which therefore will self-assemble in cells in the absence of regulatory mechanisms. The interaction between ATG9 and the HORMA domain proteins ATG13-101 is remarkably slow, but key step in the assembly of the super-complex. The conformer conversion of HORMA domains can be catalyzed, both at the assembly and the disassembly level by a specialized protein machinery (De Antoni et al., 2005; Faesen et al., 2017; Gu *et al*., 2022). This presents an enticing possibility that on-demand acceleration of the assembly or disassembly of the ATG9-13-101 complex would dynamically control contact site assembly and function in space and time. Quantifying the kinetic framework of these interactions both *in vitro* and in cells, will aid in the identification of these catalysts.

In cells, the lipid transfer protein ATG2 is recruited after the assembly of the initiation machinery. This recruitment requires the cooperative association or the activity from all canonical autophagy initiation subcomplexes: ATG13-101 from the ULK1 complex, ATG9, and the PI3-kinase complex for creating the PI3P lipid signature. This predicts that ATG2 serves as a ‘co-incidence sensor’ for the successful assembly of the super-complex. This assembled ‘minimal unit’ might then be repeated multiple times to give rise to the ‘regional’ contact sites that supports autophagosome growth. The incorporation of ATG2-WIPI4 in a larger protein network is not mutually exclusive with a model where additional ATG2 only transiently associates and subsequently diffuses onto the neighboring membranes to create a regional tether. This way the co-incidence of early autophagy initiation factors is ‘sensed’ as a mature contact site before committing to autophagosome formation.

ATG13 and ATG101 continue the trend that most HORMA domain proteins can reversibly change their protein’s three-dimensional structure to regulate their protein-protein interaction potential. However, in contrast to other HORMA domain proteins, neither ATG13 nor ATG101 require their seatbelt to interact with their interaction partners. Nevertheless, removing the seatbelt in ATG13 does change its default conformer state, suggesting it is used allosterically to create interaction interfaces. Disassembling HORMA domain effector complexes requires a significant amount of energy. This is achieved by remodelling the seatbelt by AAA^+^-ATPase TRIP13/Pch2 in an ATP-dependent manner, which ‘opens’ the seatbelt leading to the spontaneous disassembly of the effector complex (Ye et al., 2015). TRIP13 is a conserved generic HORMA remodeller, but it has not been linked to autophagy. It is therefore currently unclear if the disassembly of the ATG9-13-101 complex, and by extension the super-complexes, require ATP-dependent remodeling of the HORMA domains to silence autophagosome biogenesis.

In contrast to ATG13, ATG101 is not universally conserved and is most notably missing in *S. cerevisiae*. ATG13 binding to the N-terminus of ATG9 is conserved in *S. cerevisiae* (Suzuki *et al*.,2015), which suggests that this interaction is required for a conserved, but as-of-yet undefined function. The exact role of ATG101 is unclear. In mitosis, the ‘template model’ defines two functions for HORMA domain MAD2: a ‘template’ MAD2 acts as an enzyme to convert multiple ‘copy’ MAD2 molecules in the proper conformer to assemble the effector complex (Faesen *et al*., 2017). In human autophagy, the ‘templating’ role might be taken up by ATG101 to guide potentially multiple molecules of ATG13 to engage in the functional interaction on ATG9. Species that lack ATG101, could use the MAD2 ‘template’ mechanism more faithfully where ATG13 serves both roles. Future work will be needed to deconvolute this intricate assembly mechanism.

Detailed mechanistic studies into the architecture and assembly mechanisms of membrane contact sites are required to understand the *de novo* generation of new membrane-bound organelles, like the autophagosome. The biochemical reconstitution approach described in this work emphasizes that manipulation of recombinant initiation complexes *in vitro* allows molecular insight into crucial functions and the identification of regulatory mechanisms. Recreating the conditional assembly and disassembly of a functional contact site *in vitro* would bring us closer to the ultimate goal: the reconstitution of the initiation of autophagosome biogenesis with recombinant proteins.

## Acknowledgements

We are grateful to Anuruti Swarnkar and Claudia Schmidt and other members of the Alex Stein laboratory (MPI-NAT) for assistance with preparing liposomes and the stain-free protein quantification. We thank Stefanie Asper-Thiel for technical assistance and Faesen laboratory team members for discussion, in particular we thank Laura Griese for establishing the initial purification protocols of ULK1 and FIP200. We are grateful to Reinhard Jahn and Alex Stein for critical reading of the manuscript. This work was supported by Max Planck society (AF and HU) and DFG SFB1190 (AF)

## Author contributions

AN, FL, PH, AP, MM and ACF designed *in vitro* experiments and analyzed results. AN, FL and LG set up recombinant expression systems, established purification protocols and purified proteins with help of AP. AN, FL and AP performed *in vitro* experiments. CD and AF performed cellular assays, supervised by BS. AN and PH performed vesicle tethering, lipid scramblase and transfer experiments with help from BK, supervised by MM and ACF. IP performed crosslink mass-spectrometry and analysis, supervised by HU. ACF wrote the manuscript with input from AN and BS.

## Competing interests

Authors declare that they have no competing interests.

## Notes

### Competing Interest Statement

The authors have declared no competing interest.

### Summary of Updates

Fixed an unfortunate typo.

