## Supplemental Material for "Metamorphic proteins at the basis of human autophagy initiation and lipid transfer"

### Supplemental Information

### Reagents & antibodies

All phospholipids used in this study were purchased from Avanti Polar Lipids. Bafilomycin $A_1$ (#B1793) was purchased from Sigma-Aldrich (St. Louis, MO, USA). Additionally, the following reagents were used: Protein A and Protein G Sepharose™ 4 Fast Flow (Cytiva, #17528001 and #17061801) and HA-Agarose (Sigma-Aldrich, #A2095). For immunoblotting, primary antibodies against ß-Actin (clone AC-74, Sigma-Aldrich, #A5316), GAPDH (abcam, #ab8245), Vinculin (Sigma-Aldrich, #V9131), ATG101 (CST, #13492), ATG13 (Sigma-Aldrich, SAB4200100), ULK1 (clone D8H5, CST, #8054), FIP200 (Proteintech, #17250-1-AP), ATG14 phospho S29 (CST, #92340), ULK1 phospho S638 (CST, #14205; corresponds to murine S637), ULK1 phospho S757 (CST, #6888; corresponds to human S758), HA (Biolegend, #901501 and Roche, #11867423001), LC3B (CST, #2775), SQSTM1/p62 (PROGEN, #GP62-C), anti-mouse IgG (Santa-Cruz, #sc-2025) and anti-rabbit IgG (Merck Millipore, #12-370) were used. IRDye 680- or IRDye 800-conjugated secondary antibodies (#926-68077, #926-68076, #926-68072, #926-32212, #926-32213) were purchased from LI-COR Biosciences.

### Expression and purification of proteins

ATG9A and its transmembrane containing constructs. ATG9A with tandem N-terminal MBP and 6xHis-tag was expressed in *Trichoplusia ni* High Five™ (Hi5) insect cells using the biGBac expression system as described previously (Altmannova et al., 2021; Weissmann et al., 2016). Cell pellet was resuspended in buffer A (50 mM Hepes (pH 8), 150 mM NaCl, 5 mM $MgCl_2$, 5% (v/v) Glycerol, 0.5 mM TCEP, 1% (w/v) DDM, 0.5 mM PMSF, 10 U/ml Benzonase) as lysis buffer. Cells were lysed gently on ice for at least 1 h by a magnetic stirrer. The lysate was then diluted 3 times with DDM-free buffer A and mixed gently for another 30 mins before clarifying by centrifugation at 20,000 × *g* for 30 mins. The proteins in the supernatant were purified by affinity chromatography using MBP trap columns (Cytiva) with Buffer B (50 mM Hepes (pH 8), 150 mM NaCl, 0.5 mM TCEP, 0.03% (w/v) DDM) as binding buffer and buffer B supplement with 20 mM Maltose as elution buffer. The affinity tags were cleaved by PreScission protease treatment for at least 5 hours at 4°C after affinity chromatography (if required), followed by size exclusion chromatography using Superose 6 column (Cytiva) pre-equilibrated with Buffer B. The purified protein was concentrated, flash frozen in liquid nitrogen and stored at -80°C.

ATG9-13-101. To purify a complex of human ATG9A, ATG13, ATG101, each protein with tandem N-terminal MBP and 6xHis-tag was co-expressed in High Five™ (Hi5) cells and purified with the same protocol used for ATG9A.

$ATG9^N$. $ATG9^N$ was produced as GST or His-mC fusion constructs from Hi5 insect cells as described previously. Cells were washed and resuspended in lysis buffer C (50 mM Hepes (pH 8), 100 mM NaCl, 0.5 mM TCEP, 1 mM PMSF, 1% (v/v) Triton X-100). Resuspended cells were lysed by stirring for 1 h in the presence of DNAse I for the last 20 minutes before clearance at 15,000 × *g* at 4 °C for 30 minutes. Cleared lysate was filtered and passed over GSH-Sepharose 5-ml column before extensive washing with lysis buffer (without PMSF and Triton X-100). $ATG9^N$ was then eluted in buffer C supplemented with 20 mM reduced glutathione or 0.5 mM Maltose depending on the column used. Eluted protein was concentrated in a 50 kDa Amicon-Ultra-15 Centrifugal Filter (Millipore). Concentrated protein was then loaded onto a Superdex 200 16/600 or 10/300 GL column or a Superose 6 16/600 or 10/300 GL pre-equilibrated in Buffer D (50 mM HEPES (pH 8), 150 mM NaCl, 0.5 mM TCEP). Peak fractions corresponding to $ATG9^N$ were collected and again concentrated in a 50 kDa MWCO concentrator to at least 1 mg/ml before being flash-frozen in liquid N2 and stored at −80 °C.

$ATG9^N$-$ATG13^{HORMA}$. The $ATG9^N$-$ATG13^{HORMA}$ complex was produced as GST-$ATG13^{HORMA}$ or His-mC-$ATG9^N$ fusion constructs from Hi5 insect cells using the biGBac expression system. Bacmid was then produced as previously described for ATG101 and subsequently inserted in a pBig vector to ensure stable co-expression. Cells were washed and resuspended in lysis buffer C. Resuspended cells were lysed by stirring for 4 h in the presence of DNAse I for the last 20 minutes before clearance at 15,000 × *g* at 4 °C for 30 minutes. Cleared lysate was filtered and passed over GSH-Sepharose coupled with an HisTrap™ excel (to catch the excess unbound ATG9$^{N}$) 5-ml column before extensive washing with lysis buffer C (without PMSF and Triton X-100). ATG13 was then eluted in lysis buffer supplemented with 20 mM reduced glutathione or 150 mM imidazole-containing elution buffer depending on the column used. Eluted protein was concentrated in a 50 kDa Amicon-Ultra-15 Centrifugal Filter (Millipore) and then loaded onto a Superose 6 16/600 or 10/300 GL pre-equilibrated in buffer D. Peak fractions corresponding to ATG13-ATG9$^{N}$ were collected and again concentrated in a 50 kDa MWCO concentrator to at least 1 mg/ml before being flash-frozen in liquid N2 and stored at −80 °C.

ATG9$^{C}$. ATG9$^{C}$ and its mutant were produced as GST fusion constructs from *E. coli* expression system (Rosetta and Lobster expression cell lines). Cells were washed and resuspended in lysis buffer E (50 mM Hepes (pH 8), 100 mM NaCl, 0.5 mM TCEP, 1 mM PMSF, 1% Triton X-100, and lysozyme) and lysed by stirring for 1 h before clearance at 15,000 × *g* at 4 °C for 30 minutes. Cleared lysate was passed over GSH-Sepharose 5-ml column before extensive washing with lysis buffer (without PMSF and Triton X-100). ATG9$^{N}$ was then eluted in lysis buffer + 20 mM reduced glutathione. Eluted protein was concentrated in a 10 kDa Amicon-Ultra-15 Centrifugal Filter (Millipore). Concentrated protein was then loaded onto a Superdex 200 16/600 or 10/300 GL column pre-equilibrated in buffer D. Peak fractions corresponding to ATG9$^{C}$ were collected and again concentrated in a 10 kDa MWCO concentrator to at least 1 mg/ml before being flash-frozen in liquid N2 and stored at −80 °C.

ATG13. ATG13$^{HORMA}$ and its mutants were produced as GST or His-MBP fusion constructs from Hi5 insect cells using the biGBac expression system. Cells were washed and resuspended in lysis buffer C. Resuspended cells were lysed by stirring for 4 h in the presence of DNAse I for the last 20 minutes before clearance at 15,000 × *g* at 4 °C for 30 minutes. Cleared lysate was filtered and passed over GSH-Sepharose or MBPTrap™ HP 5-ml column before extensive washing with lysis buffer (without PMSF and Triton X-100). ATG13 was then eluted in lysis buffer C + 20 mM reduced glutathione or 0.5 mM Maltose depending on the column used. Eluted protein was concentrated in a 50 kDa Amicon-Ultra-15 Centrifugal Filter (Millipore). Concentrated protein was then loaded onto a Superdex 200 16/600 or 10/300 GL column or a Superose 6 16/600 or 10/300 GL pre-equilibrated in buffer D. Peak fractions corresponding to ATG13 were collected and again concentrated in a 50 kDa MWCO concentrator to at least 1 mg/ml before being flash-frozen in liquid N2 and stored at −80 °C.

Full-length ATG13 (His-MBP-tagged) was expressed in Hi5 insect-cells. Cell pellets were resuspended in 50 ml lysis buffer F (25 mM Hepes (pH 8), 300 mM NaCl, 5% (v/v) Glycerol, 0.5 mM TCEP, 1 mM EDTA, protease inhibitor cocktail) and sonicated for 5 min on-time by pulsing 5 s on/ 10 s off at an amplitude of 25%. The lysate was then stirred magnetically after addition of 0.2% DDM, 10 µg/ml DNAse, 5 mM $MgCl_2$ for 30 min, after which it was diluted to 250 ml with dilution buffer G (25 mM Hepes (pH 8.0), 300 mM NaCl, 5% (v/v) Glycerol, 0.5 mM TCEP). Clarification of lysate was done by centrifugation at 10,000 rpm for 30 min at 4 °C and supernatant loaded on a 5 ml MBPTrap column (Cytiva). Column was equilibrated for 5 CV with wash buffer H (25 mM HEPES (pH 8.0), 300 mM NaCl, 5% Glycerol, 0.5 mM TCEP) and elution was done by addition of 5 mM Maltose in the wash buffer. Following affinity purification, peak fractions were pooled, concentrated with a centrifugal concentrator (MWCO 50 kDa, Amicon, MERCK), measured for amount of protein and loaded on a Superose 6 16/600 HiLoad column equilibrated with buffer I (25 mM HEPES, pH 8.0, 300 mM NaCl, 5% (v/v) Glycerol, 0.5 mM TCEP). For cleaving the 6X-His-MBP tag, 1:1000 PreScission Protease was added to the concentrated fractions and applied for SEC. After analysis by SDS-PAGE, fractions corresponding to ATG13 were concentrated and stored by flash-freezing at -80 °C.

ATG101. ATG101 was produced as GST, His-MBP, or Strep fusion constructs from Hi5 insect cells using the biGBac expression system. Cell pellets were washed and resuspended in lysis buffer C. Resuspended cells were lysed by stirring for 1 h in the presence of DNAse I for the last 20 minutes before clearance at 15,000 × *g* at 4 °C for 25 minutes. Cleared lysate was filtered and passed over GSH-Sepharose, MBPTrap™ HP, or *Strep*-Tactin Superflow 5-ml column before extensive washing with lysis buffer C (without PMSF and Triton X-100). ATG101 was then eluted in lysis buffer + 20 mM reduced glutathione, 0.5 mM Maltose, or 0.5 mM DSB depending on the column used. Eluted protein was concentrated in a 30 kDa or 10 kDa Amicon-Ultra-15 Centrifugal Filter (Millipore) in the presence or absence of GST- or His-tagged 3C protease (incubation over-night at 4 °C). Concentrated protein was then loaded onto a Superdex 200 16/600 or 10/300 GL column or Superdex 75 16/600 or 10/300 GL column pre-equilibrated in buffer D. In the case of the cut protein, a 1 ml GSH-Sepharose or a HisTrap™ excel was connected in series after the Superdex column to trap the tag, un-cut ATG101, and tagged 3C protease. Peak fractions corresponding to ATG101 were collected and again concentrated in a 30 or 10 kDa MWCO concentrator to at least 1 mg/ml before being flash-frozen in liquid N2 and stored at −80 °C.

ATG2A. Human ATG2A with a C-terminal StrepII tag expressed in High Five™ (Hi5) cells were purified similarly by affinity chromatography using Strep trap columns (Cytiva) with buffer A (50 mM Hepes (pH 8), 150 mM NaCl, 5 mM $MgCl_2$, 5% (v/v) Glycerol, 0.5 mM TCEP, 1% (w/v) DDM, 0.5 mM PMSF, 10 U/ml Benzonase) as lysis buffer, Buffer J (50 mM Hepes (pH 8), 150 mM NaCl, 0.5 mM TCEP, 0.1% (w/v) DDM) as binding buffer, and buffer J supplement with 5 mM desthiobiotin as elution buffer. The proteins were finally purified by size exclusion chromatography using Superose 6 column (Cytiva) pre-equilibrated with Buffer C. The purified protein was concentrated, flash frozen in liquid nitrogen and stored at -80°C. For lipid transfer assay, DDM was subsequently removed by two successive incubations with Pierce detergent removal resin (Thermo Fisher Scientific), pre-equilibrated with DDM-free buffer C. Incubation in the detergent removal resin was performed at room temperature for 15 min each.

ULK1. ULK1 with N-terminal GST tag was expressed in High Five™ (Hi5) cells and purified similarly by affinity chromatography using GST trap column (Cytiva). Cells were lysed gently in buffer K (50 mM Hepes (pH 8), 500 mM NaCl, 5 mM $MgCl_2$, 5% (v/v) Glycerol, 0.5 mM TCEP, 1% (w/v) DDM, 0.5 mM PMSF, 10 U/ml Benzonase, 100 μM Leupeptin) on ice for 1 h by a magnetic stirrer. The lysate was then diluted 3 times with DDM-free buffer K and mixed gently for another 30 mins. The protein was purified by affinity chromatography using GST trap column (Cytiva) with binding buffer L (50 mM Hepes (pH 8), 500 mM NaCl, 5% (v/v) Glycerol, 0.5 mM TCEP, 0.5 mM EDTA, 0.03% (w/v) DDM, 0.5 mM PMSF, 100 μM Leupeptin), and buffer L supplement with 10 mM reduced glutathione as elution buffer. The protein was subjected to size exclusion chromatography with Superose 6 column (Cytiva) pre-equilibrated with Buffer L. The purified protein was concentrated, flash frozen in liquid nitrogen and stored at -80°C.

FIP200. FIP200 with tandem N-terminal MBP and 6xHis-tag expressed in High Five™ (Hi5) cells was purified similarly by affinity chromatography using MBP trap columns (Cytiva) with buffer M (50 mM Hepes (pH 8), 500 mM NaCl, 5 mM $MgCl_2$, 5% (v/v) Glycerol, 0.5 mM TCEP, 0.5 mM EDTA, 1% (w/v) DDM, 0.5 mM PMSF, 10 U/ml Benzonase) as lysis buffer, Buffer N (50 mM Hepes (pH 8), 500 mM NaCl, 5% (v/v) Glycerol, 0.5 mM TCEP, 0.5 mM EDTA, 0.03% (w/v) DDM, 0.5 mM PMSF) as binding buffer, and buffer N supplemented with 20 mM Maltose as elution buffer. The proteins were finally purified by size exclusion chromatography using Superose 6 column (Cytiva) pre-equilibrated with Buffer E. The purified proteins were concentrated, flash frozen in liquid nitrogen and stored at -80°C.

ATG14-BECN1. The ATG14 and BECN1 complex was produced as His-GFP fusion ATG14 construct and BECN1 from High Five™ (Hi5) insect cells using the biGBac expression system. Cell pellets were washed and resuspended in lysis buffer O (50 mM Hepes (pH 8), 200 mM NaCl, 5% (v/v) Glycerol, 5 mM Imidazole, 0.5 mM TCEP, 1 mM PMSF, 1%Triton X-100). Resuspended cells were lysed by stirring for 2 h in the presence of DNAseI for the last 20 minutes before clearance at 15,000 x *g* at 4°C for 30 minutes. Cleared lysate was filtered and passed over a HisTrap™ excel 5 ml column before extensive washing with lysis buffer O (without PMSF and Triton X-100). ATG14-BECN1 complex was then eluted with the lysis buffer containing 300 mM Imidazole through an elution gradient. The complex eluted around 30% elution buffer was then incubated overnight at 4°C in the presence of His-tagged 3C protease and subsequently loaded onto a HiQ column to separate the tag using a salt gradient. The complex eluted as a complex around 400 mM NaCl. Eluted product was concentrated in a 100kDa

Amicon-Ultra-15 Centrifugal Filter (Millipore) and loaded on a Superdex 200 16/600 or 10/300 column pre-equilibrated in buffer P (50 mM HEPES (pH 8), 300 mM NaCl, 0.5 mM TCEP). Peak fractions corresponding to ATG14-BECN1 were collected and again concentrated in a 100 kDa MWCO concentrator to at least 1 mg/ml before being flash-frozen in liquid N2 and stored at −80 °C.

ATG14 with tandem N-terminal His-mCherry tag and was co-expressed with BECN1 with a tandem N-terminal His-MBP tag in Hi5 insect cells using the biGBac expression system. Cell pellets were harvested and purified using the same protocol used for FIP200.

WIPI4. WIPI4 with different tag variants (N-terminal GST tag, tandem N-terminal GFP tag and 6xHis-tag, tandem N-terminal mC tag and 6xHis-tag) were expressed in High Five™ (Hi5) cells and purified similarly by affinity chromatography using appropriate affinity column. Cells pellets were lysed by sonication in DDM-free buffer A. Clarified lysate was applied to appropriate affinity chromatography columns using DDM-free buffer A as binding buffer and eluted by either 0 to 300 mM Imidazole gradient or 10 mM reduced glutathione depending on the column used. The tag was cleaved if required as described above and the proteins were finally purified by size exclusion chromatography using Superdex S200 column (Cytiva) pre-equilibrated with binding buffer. The purified protein was concentrated, flash frozen in liquid $N_2$ and stored at -80°C.

**Analytical size-exclusion chromatography (SEC) analysis.**

Analytical SEC analysis was performed using the indicated column in SEC buffer containing 250 mM NaCl, 25 mM HEPES (pH 8), 0.5 mM TCEP, 2.5% Glycerol on an Äkta Pure system. All samples were eluted under isocratic conditions at 4°C in SEC buffer at a flow rate of 0.5 ml/min. Elution of proteins was monitored by absorption of light at 280 nm. Fractions (0.5 ml) were collected and analyzed by SDS-PAGE and Coomassie blue stain. To detect the formation of a complex, proteins were mixed at the indicated concentrations in 250 µl, incubated on ice for at least 1 h and then subjected to SEC.

**GST and MBP pulldown assays**

GST, MBP and Strep pulldown experiments were performed using pre-blocked GSH Sepharose beads, MBP beads (Amylose resin) (NEB), or StrepTactin Superlow Plus (Qiagen) beads in respective pulldown buffers. The pulldowns, which required incubation of the baits and prey for less than 1 h, were performed by pre-binding the bait to 30 µl of beads before the prey was added. In case of long incubation from 1 h to 24 h, the prey and bait were pre-formed and then loaded on the beads for 10 minutes. At the same time, 1 µg of each protein was added into Laemmli sample loading buffer for the input gel. In pulldowns containing ULK1, we used the ULK1 lysis buffer (buffer L) during incubation and reduced DDM concentration to 0.03% during washing. Beads were spun down at 500 × *g* for 1 min. The supernatant was removed, and beads were washed at least twice with 500 µl buffer. The supernatant was removed completely, samples boiled in 10 µl Laemmli sample loading buffer, and run on a 12% SDS–PAGE gel. Bands were visualized with Coomassie brilliant blue staining. For information regarding concentration, temperature, and variations, see figure legends.

**Stain-free protein quantification**

Stain-Free (SF) is a method of protein visualization and quantification which enables detection of protein bands in gels without using colorimetric or fluorescent stains (Gurtler et al., 2013; Holzmuller and Kulozik, 2016). SF gels contain a trihalo compound within the gel matrix which reacts with tryptophan residues using an ultraviolet light-induced reaction to produce fluorescent light. The fluorescence allows visualization of proteins in the gel without additional staining and destaining steps. In this study, to quantify the relative amount of each protein in a complex, purified protein complexes in increasing amount (5 µg, 10 µg, 15 µg, 20 µg) were loaded on 12% Mini-PROTEAN® TGX Stain-

Free™ Protein Gels (Bio-Rad Laboratories) and separated by electrophoresis. After electrophoresis protein bands were visualized by placing the gel on the UV transilluminator Gel Doc™ EZ imager (Bio-Rad Laboratories). Intensities of protein bands were then normalized against the number of Tryptophan residue in each protein. A linear fit of the band intensities against the amount of protein was fitted to verify that the intensity proportionally increases with the increase in protein loaded (5 µg, 10 µg, 15 µg, 20 µg).

**Preparation of protein-free (fluorescent donor and acceptor) liposomes**

Large Unilamellar Vesicles (LUVs) were prepared by reversed-phase evaporation as described previously *(Hernandez et al., 2012)*. Briefly, lipids were dissolved in chloroform and mixed at a desired molar ratio (donor LUVs: 46% DOPC, 25% DOPE, 20% DOPS, 2% NBD-PE, and 2% Rh-PE, 5% PI3P; acceptor LUVs: 50% DOPC, 25% DOPE, and 25% DOPS). Chloroform was subsequently removed using a rotary evaporator to allow lipid film formation. The lipid film was then dissolved in 1 ml diethyl ether, followed by 300 µl of buffer F (50 mM Hepes (pH 8), 150 mM NaCl, 0.5 mM TCEP). The sample was then sonicated for 1 min in a bath sonicator at 4°C to create emulsion. Diethyl ether was initially removed at 500 mbar for 10 min, and 700 µl of buffer F was added. The remaining diethyl ether was removed by lowering the pressure stepwise to 100 mbar until diethyl ether was completely removed. The resulting lipid suspension was extruded 11 times through a 0.4-µm polycarbonate filter and then 21 times through a 0.1-µm polycarbonate filter (Mini extruder kit, Avanti Polar Lipids).

**Reconstitution of proteins into liposomes**

To reconstitute ATG9, protein-free LUVs was destabilized by the addition of DM at the concentration described by the R-value (Rigaud and Levy, 2003). ATG9 was added at a protein:lipid ratio of 1:2000 or 1:500 depending on the experiments. The solution was incubated for 1 h at room temperature and DM was subsequently removed by three successive incubations with Pierce detergent removal resin (Thermo Fisher Scientific), pre-equilibrated with buffer F. Incubation in the detergent removal resin was performed at room temperature for 15 min each.

**Flotation of reconstituted liposome**

Flotation assay was performed as described previously (Krick et al., 2012) to verify proper reconstitution of ATG9 into protein-free liposome. Briefly, 50 µl of ATG9 proteoliposomes were mixed with 50 µl 80% (w/v) Nycodenz (Alere Technologies) prepared in buffer F. The mixture was subsequently overlaid with 40 µl of 30% Nycodenz, 40 µl 15% Nycodenz and 40 µl of buffer F, respectively. The density gradient was centrifuged at 50,000 rpm in a S55-S swinging bucket rotor (Thermo Fisher Scientific) for 1 h at 4 °C. Six equal fractions was collected from the top of the gradient and analyzed by SDS-PAGE.

**Protease protection assay**

The orientation of ATG9-MBP in liposomes was determined by assessing the accessibility of the N-terminal HRV 3C-cleavage site to PreScission protease. Proteoliposomes were incubated with 10 µM PreScission protease at 4°C for 20 mins, 40 mins, or overnight. In the control, 1% DDM was added to the proteoliposome and precision protease mix. The reactions were stopped by addition of SDS loading buffer and samples were analyzed by SDS-PAGE. Gels were quantified using Fiji (ImageJ) (Schindelin et al., 2012).

**Lipid transfer assay**

To monitor protein mediated lipid transfer between liposomes, we performed FRET-based dequenching assays as described previously (Connerth et al., 2012; Miyata et al., 2016; Watanabe et al., 2015). In brief, a mixture of donor liposomes containing both NBD-PE and fluorescent rhodamine (Rhod)–labeled PE (46% DOPC, 25% DOPE, 20% DOPS, 2% NBD-PE, and 2% Rh-PE, 5% PI3P), and acceptor liposomes without fluorescent lipids were prepared (50% DOPC, 25% DOPE, and 25% DOPS). In donor liposome, NBD-PE fluorescence is quenched by Rhod-PE due to their proximity. If lipid exchange between liposomes happens when lipid transfer proteins are introduced into the mixture, the NBD signal will be dequenched and increase in intensity. In our experiments, 25 μM lipid concentration of donor liposomes and acceptor liposomes in the presence or absence of the indicated proteins except ATG2 was prepared in 200 μl buffer A on a 96-well microplate (Greiner bio-one). In the sample containing ATG9, the protein was reconstituted into both donor and acceptor liposomes. The microplate was placed in a Synergy Neo 2 Multi-Mode Reader (BioTek) and gently shaken for 30 mins at 25°C. Subsequently, ATG2 was added at the desired concentrations to start the reaction and NBD fluorescence intensity (excitation, 485 nm; emission, 528 nm) was monitored for 2 or 3 hours. A lag time was expected between samples due to preparation of the mixtures. After the indicated time, Triton X-100 was added to the reaction mixture at 0.5% (v/v) final concentration to solubilize all lipids and therefore maximize NBD fluorescence signal, the signal was then monitored until stabilized. The activities shown in one figure were simultaneously measured. All proteins used in lipid transfer were un-tagged except ATG2 with C-terminal StrepII tag. All data were normalized as a percentage of total NBD fluorescence after Triton-X100 addition. Transfer rate ($k_{obs}$) was obtained by fitting the data to a one-phase exponential association equation using GraphPad Prism.

**Scramblase assay**

The scramblase assay of ATG9 was performed as previously reported (Menon et al., 2011; Ploier and Menon, 2016). In brief, proteo-liposome or protein-free liposome containing a trace amount of NBD-PE distributed equally between both leaflets are supplemented with dithionite which irreversibly quenches NBD fluorophore on the outer leaflet, but not the inner leaflet due to its impermeable property. Hence, dithionite addition leads to a 50% decrease of fluorescence for protein free liposomes and a greater reduction for scramblase-containing proteo-liposomes due to the rapid exchange of lipids between leaflets induced by the scramblase protein. In our experiments, 50 μM of protein-free liposome/ATG9 proteo-liposome-containing NBD-lipids were prepared in 200 μl buffer A on a 96-well microplate (Greiner bio-one). The microplate was placed in a Synergy Neo 2 Multi-Mode Reader (BioTek) and NBD fluorescence intensity (excitation, 485 nm; emission, 528 nm) was monitored. After initial signal stabilization, the solution was supplemented with 50 mM sodium dithionite and further supplemented with 50 mM dithionite and Triton X-100 0.5% (v/v) after about 600 s of incubation.

**CXL-MS analysis**

For ATG9-ATG13H-ATG101 protein–protein crosslinking, 10 μg aliquots of the complex were incubated with either 0.25, 0.5 or 1 mM BS3 (Thermo Scientific). Each of the samples was incubated at 4°C for 60 mins and subsequently quenched by a final concentration of 50 mM Tris for 15 mins. Proteins were then separated by PAGE using a 4–12% gradient gel (NuPAGE, Invitrogen). The cross-linked complex was cut out of the gel. Excised gel pieces were then subjected to in-gel tryptic digest (Shevchenko et al., 2006). Samples were reduced with 10 mM dithiotreitol and alkylated with 55 mM iodacetamide and subsequently digested with trypsin (sequencing grade, Promega) at 37 °C for 18 h. Extracted peptides were dried in a SpeedVac Concentrator and dissolved in loading buffer composed of 4% acetonitrile and 0.05% TFA. Samples were subjected to liquid chromatography mass spectrometry (LC-MS) on a QExactive HF-X (Thermo Scientific). Peptides were loaded onto a Dionex UltiMate 3000 UHPLC+ focused system (Thermo Scientific) equipped with an analytical column (75 µm x 300 mm, ReproSil-Pur 120 C18-AQ, 1.9 µm, Dr. Maisch GmbH, packed in house). Separation by reverse-phase chromatography was done on a 60 min multi-step gradient with a flow rate of 0.3-0.4 µl $min^{-1}$. MS1 spectra were recorded in profile mode with a resolution of 120 k, maximal injection time was set to 50 ms and AGC target to $1e^6$ to acquire a full MS scan between 380 and 1580 m/z. The top 30 abundant precursor ions (charge state 3-8) were triggered for HCD fragmentation (30% NCE). MS2 spectra were recorded in profile mode with a resolution of 30 k; maximal injection time was set to 128 ms, AGC target to $2e^5$, isolation window to 1.4 m/z and dynamic exclusion was set to 30 s. Raw files were analyzed via pLink2.3.5 to identify cross-linked peptides (Chang et al., 2015). Database was generated based on the protein complex used. FDR was set to 1% and results were filtered by excluding crosslinks supported by only one cross-linked peptide spectrum match. The crosslinks were visualized using the webserver xiNET (Combe et al., 2015).

**Fluorescence microscopy of liposomes**

For Microscopy the LSM 780 (Carl Zeiss) was used. µ-slides with 8-wells (Ibidi)were coated with BSA by incubating each well with 100 µl 5 mg/ml BSA followed by 3 washing cycles with protein buffer (150 mM NaCl, 50 mM HEPES, 0.5 mM TCEP,1 mM EDTA, pH 8, Osm. 380 mOsm). The wells were prepared for the addition of the GUVs with 200 µl of protein buffer. GUVs were added carefully with a tip-cut pipette. An appropriate window for microscopy was selected and indicated proteins were added to the wells. The pictures were processed with ImageJ-software.

**Generation of GUVs**

GUVs were formed by an adapted electroformation protocol as described before (Kroppen et al., 2021; Tarasenko et al., 2017) in the VesiclePrepPro (Nanion). In brief, first a lipid mix with the end concentration of 2 mg/ml was prepared. A rubber ring (Ø28 mm) was slightly coated with silicon and placed carefully on the center of the electrically conductive side of an ITO-plate. The ITO-plate was heated to 50°C on a heating plate. 7.5 µl of the lipid mixture was applied dropwise with a Hamilton syringe on the ITO-plate in the area surrounded by the rubber ring. Following the ITO-plate was placed in a vacuume chamber for 10 min to evaporate the residual organic solvent. The plate was inserted in the chamber and an electroformation buffer (240 mM sucrcose, 50 mM HEPES, pH 8, Osm. 380 mOsmol) was added slowly on the lipids. A second ITO-plate was placed on top of the first ITO-plate with the electrically conductive side facing the lipids and the buffer. This way the chamber was sealed.

The electroformation protocol used here consists of three phases: In phase 1 the peak-to-peak amplitude rises linearly from 0 to 2 V. During phase 2 it stays on 2 V for 2 h 55 min. In phase 3 the amplitude decreases to 0 V again in a 20 min period. The frequency is set to 10 Hz in phase 1 and 2. In phase 3 it decreases to 0 Hz linearly. The temperature is set across all three phases to 55 °C and as such above the phase-transition temperature of the lipid mix. After finishing the protocol, GUVs were harvested into 1.5 ml Eppendorf tubes. They were used immediately for microscopy.

**DLS analysis**

LUVs were prepared as described above. To mimic conditions of the lipid transfer experiment, 25 µM lipid concentration of donor liposomes and acceptor liposomes in the presence or absence of the indicated proteins except ATG2 were prepared in 200 ul buffer. A 72-well Terasaki-plate (Greiner bio-one), was prepared with a thin layer of liquid parafinol. 1 µl of each solution was added to the wells. The plate was measured with the spectralight 610 on automatic settings. For creating the diagrams,

GraphPad Prism (GraphPad Software, Inc.) software was used.

**Cell lines and cell culture**

Wild-type and *Atg13* KO MEFs containing an insertion of a gene-trap cassette in the *Atg13* gene, and wild-type and *Atg101* KO MEFs were kindly provided by Noboru Mizushima (Department of Biochemistry and Molecular Biology, Graduate School and Faculty of Medicine, University of Tokyo, Japan) and have previously been described (Kaizuka and Mizushima, 2015; Suzuki et al., 2015a). Wild-type MEFs, *Atg13* KO MEFs, *Atg101* KO MEFs and corresponding transfectants generated in this study were cultured in high-glucose (4,5 g/L) DMEM (Gibco, #41965-039) supplemented with 10% FCS (Sigma-Aldrich, #F0804, LOT BCCB7649), 100 U/ml penicillin and 100 µg/ml streptomycin (Gibco, #15140-122) at 37 °C and 5% $CO_2$ humidified atmosphere. For amino acid starvation, cells were washed once with DPBS (Dulbecco's Phosphate-Buffered Saline, Gibco, #14190-094) and incubated for the indicated time points in EBSS (Earle's Balanced Salt Solution, Gibco, #24010-043).

**Retroviral transduction**

Generation of pMSCVpuro-HA-ATG13 has previously been described (Hieke et al., 2015) pMXs-IP-3xHA-ATG101 was kindly provided by Noboru Mizushima (Department of Biochemistry and Molecular Biology, Graduate School and Faculty of Medicine, University of Tokyo, Japan). For generation of pMXs-IP empty vector control the sequence encoding 3xHA-ATG101 was excised by *NotI* (Thermo Fisher Scientific, #FD0595) digestion and the backbone was subsequently ligated using T4 ligase (Thermo Fisher Scientific, #EL0011). For generation of cDNAs encoding HA-ATG13_ΔSB2 or 3xHA-ATG101_ΔN12, site-directed mutagenesis was performed using the following primers: ΔSB2 fwd: TTCATGTCTACCAGGCAATTTG, ΔSB2 rev: AGTGATGGTGCCCACAGG, ΔN12 fwd: GAGGGGCGGCAGGTGGAG, ΔN12 rev: CATTCTAGAGGTACCACGCGTGAATTC. Plat-E cells (kindly provided by Toshio Kitamura, Institute of Medical Science, University of Tokyo, Japan) were used as packaging cells, and were transfected with the pMSCVpuro or pMXs-IP-based retroviral expression vectors using FuGENE® 6 (Promega, #E2691). After 48 h, MEFs were incubated with the corresponding retroviral supernatants containing 3 µg/ml Polybrene (Sigma-Aldrich, #H9268-106) and selected in medium containing 2.5 µg/ml puromycin (InvivoGen, #ant-pr-1).

**Immunoblotting**

Cells were harvested by scraping, washed once with ice-cold phosphate-buffered saline (PBS). For immunoblotting, cells were lysed in standard ice-cold lysis buffer (20 mM Tris-HCl (pH 7.5), 150 mM NaCl, 0.5 mM EDTA, 1% [v/v] Triton X-100, protease inhibitor cocktail [Roche, #58698000] and PhosSTOP [Roche, #04906837001]) for 30 min on ice. Lysates were clarified by centrifugation at 13,300 rpm for 15 min at 4 °C. Equal amounts of protein were determined by Bradford method and subjected to SDS-PAGE. Proteins were then transferred to PVDF membranes (Millipore, #IPFL00010) and analyzed using the indicated primary antibodies and appropriate IRDye-conjugated secondary antibodies. Protein signals were detected using an Odyssey Infrared Imaging system (LI-COR Biosciences) and quantified using Image Studio Lite 5.2 (LI-COR Biosciences).

**Statistics and reproducibility**

All lipid transfer and scramblase experiments were done in independent triplicates. Comparisons among different variants were determined by one-way analysis of variances (ANOVA), followed by Turkey's multiple comparisons test. Student's t-test was used for 2-group comparisons. The error bars of these experiments indicate the standard error. P values < 0.05 were considered statistically significant. For immunoblotting, the density of each protein band was divided by the average density of all bands of this protein. The ratios were normalized to the loading control, and fold changes were calculated by dividing each normalized ratio by the average of the ratios of the control line (n=3). The results are shown as mean + standard deviation. For Figure 2d and 2e and for Ext. data Figure 4a and 4b, statistical analysis was performed using ordinary two-way ANOVA (corrected by Tukey's multiple comparisons test). For Ext. data Figure 3d, statistical analysis was performed using ordinary one-way ANOVA (corrected by Dunnett's multiple comparisons test). Compared treatments or cell lines are indicated in the corresponding bar diagrams. P values < 0.05 were considered statistically significant. All statistical data were calculated with GraphPad Prism (version 9.0.0).

### Supplemental Figures and Figure Legends

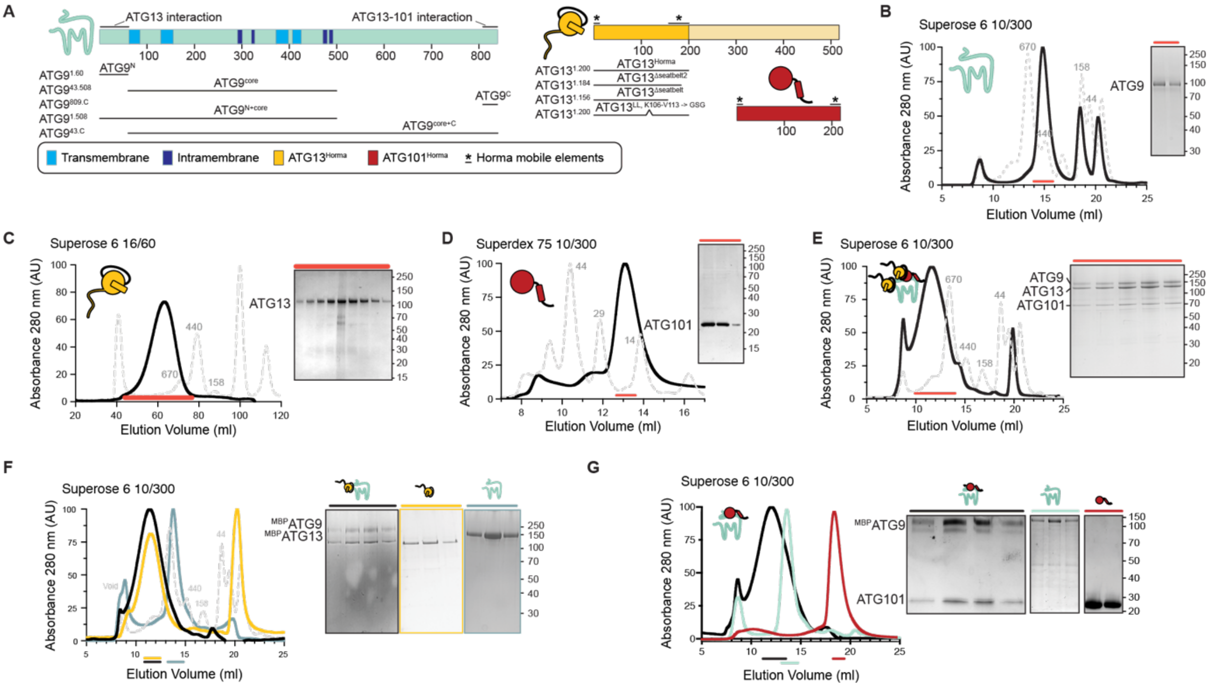


**Figure S1.** (**A**) Schematic drawing of ATG9, ATG13, ATG101 constructs used in this study. (**B**) SEC profiles of ATG9 (black trace), and marker proteins (gray dash-line trace) run on a Superose 6 10/300 SEC column and analyzed by SDS–PAGE. (**C**) SEC profiles of ATG13 (black trace), and marker proteins (gray dash-line trace) run on a Superose 6 16/600 SEC column and analyzed by SDS–PAGE. (**D**) SEC profiles of ATG101 (black trace), and marker proteins (gray dash-line trace) run on a Supedex 75 10/300 SEC column and analyzed by SDS–PAGE. (**E**) SEC profiles of $^{MBP}$ATG9-$^{MBP}$ATG13-$^{MBP}$ATG101 (black trace), and marker proteins (gray dash-line trace) run on Superose 6 10/300 SEC column and analyzed by SDS–PAGE. (**F**) SEC profiles of $^{MBP}$ATG9-$^{MBP}$ATG13 complex (black trace), $^{MBP}$ATG13 (yellow trace), $^{MBP}$ATG9 (light blue trace) and marker proteins (gray dash-line trace) run on Superose 6 10/300 SEC column and analyzed by SDS–PAGE. Analysis of corresponding peak fractions demonstrates co-elution of the subunits ($^{MBP}$ATG9 and $^{MBP}$ATG13). (**G**) SEC profiles of $^{MBP}$ATG9-$^{MBP}$ATG101 complex (black trace), $^{MBP}$ATG101 (yellow trace), $^{MBP}$ATG9 (light blue trace) and marker proteins (gray dash-line trace) run on Superose 6 10/300 SEC column and analyzed by SDS–PAGE. Analysis of corresponding peak fractions demonstrates co-elution of the subunits ($^{MBP}$ATG9 and $^{MBP}$ATG101).

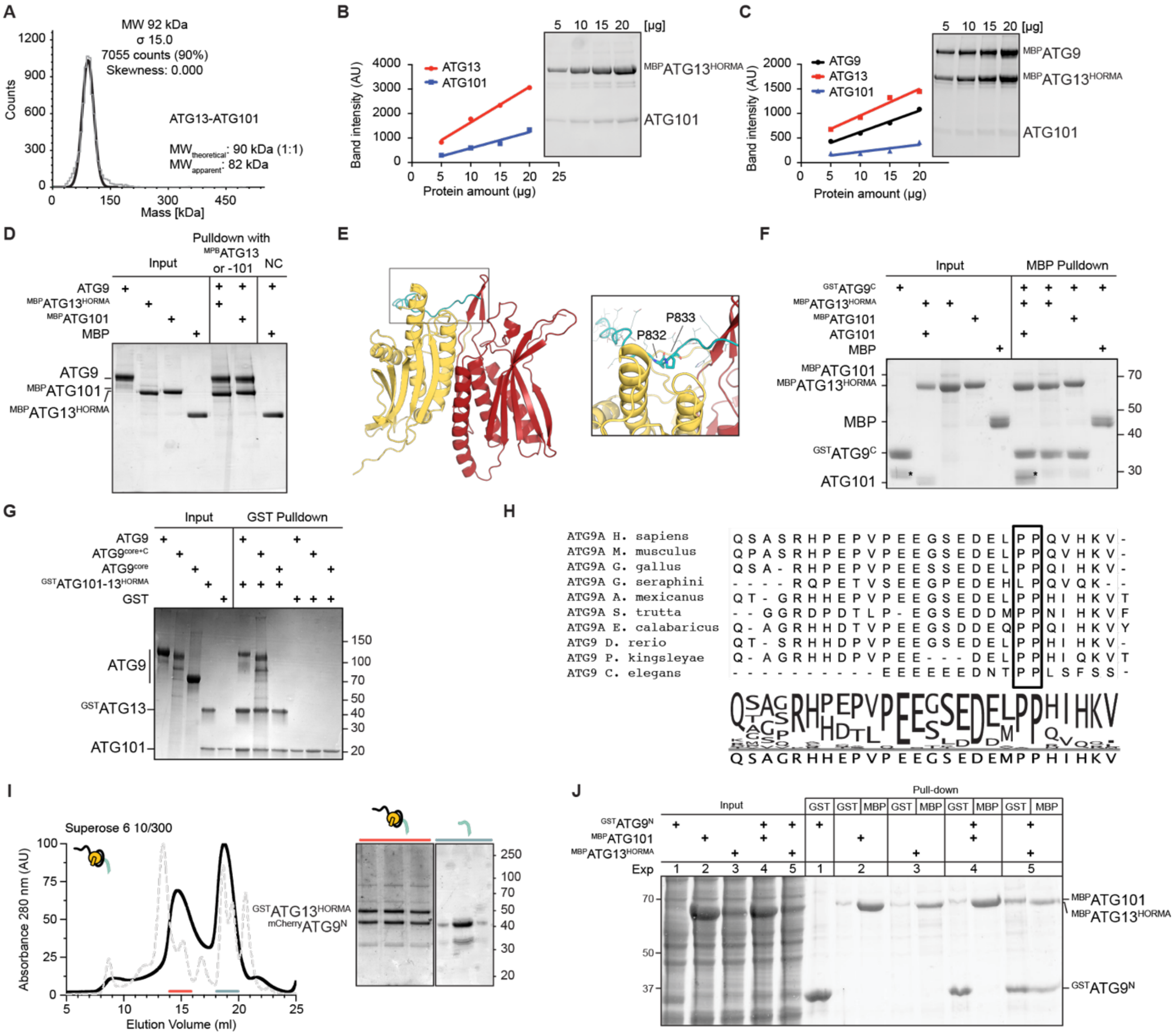


**Figure S2.** (**A**) Mass photometry profile of $^{MBP}$ATG13$^{HORMA}$-101 showing an experimental MW at 87 kDa compared to theoretical MW of a 1 to 1 stoichiometry complex at 90 kDa. (**B**) Linear dynamic assessment of the complex $^{MBP}$ATG13$^{HORMA}$-101. Indicated amounts of 1:1 stoichiometry $^{MBP}$ATG13 $^{HORMA}$-101 protein complex were analyzed on a stain-free gel. Band intensities were normalized against the number of tryptophan residues in each protein. (**C**) Indicated amount of $^{MBP}$ATG9-$^{MBP}$13$^{HORMA}$ -101 complex analyzed on a stain-free gel. Band intensities were normalized against the number of tryptophan residues in each protein. (**D**) ATG13$^{HORMA}$ and ATG101 interact with ATG9. Pull-down experiment using $^{MBP}$ATG13$^{HORMA}$, $^{MBP}$ATG101, or MBP as bait. The assay was performed with 3 µM ATG9 and 1 µM of bait. Proteins were incubated at 20 °C for 24 h. The experiment was repeated at least three times with basically identical results. (**E**) ATG9$^{829\text{-}839}$-13$^{HORMA}$-101 complex structure predicted by AlphaFold2. ATG13 is colored yellow, ATG101 is red and ATG9 is cyan. (**F**) ATG9$^{C}$ interacts with ATG13, ATG101 and the ATG13$^{HORMA}$-101 complex. Pull-down experiment using $^{GST}$ATG9$^{C}$ as prey and $^{MBP}$ATG13-101, $^{MBP}$ATG13$^{HORMA}$ and $^{MBP}$ATG101 as baits. Proteins were incubated at 20 °C for 24 h. The assay was performed with 5 µM of bait and 30 µM of prey. The experiment was repeated at least three times with basically identical results. Asterisk denotes GST degradation. (**G**) ATG13$^{HORMA}$-101 complex interacts with ATG9 and ATG9$^{C}$, but not with ATG9$^{core}$. Pull-down experiment using $^{GST\text{-}}$ATG13$^{HORMA}$-101 as bait. The assay was performed with 3 µM $^{MBP}$ATG9 truncations and 1 µM of bait. Proteins were incubated at 4 °C for 18 h. (**H**) Conservation of ATG9$^{C}$. Alignment made using HMMER (Finn et al., 2015) and ClustalOmega2 (Sievers et al., 2011). (**I**) ATG13$^{HORMA}$ binds ATG9$^{N}$. Recombinant $^{His\text{-}mCherry}$ATG9$^{N}$ and $^{GST}$ATG13$^{HORMA}$ co-elute in SEC. The shift in the ATG9$^{N}$ peak elution profile indicates complex formation. (**J**) ATG9$^{N}$ interacts with ATG13$^{HORMA}$ but not ATG101. Pull-down experiment using Hi5 lysates where $^{GST\text{-}}$ATG9$^{N}$ was expressed in isolation or in presence of $^{MBP}$ATG13$^{HORMA}$ or $^{MBP}$ATG101. Negative controls are $^{MBP}$ATG13$^{HORMA}$ and $^{MBP}$ATG101 expressed in isolation. The experiment was repeated at least three times with basically identical results.

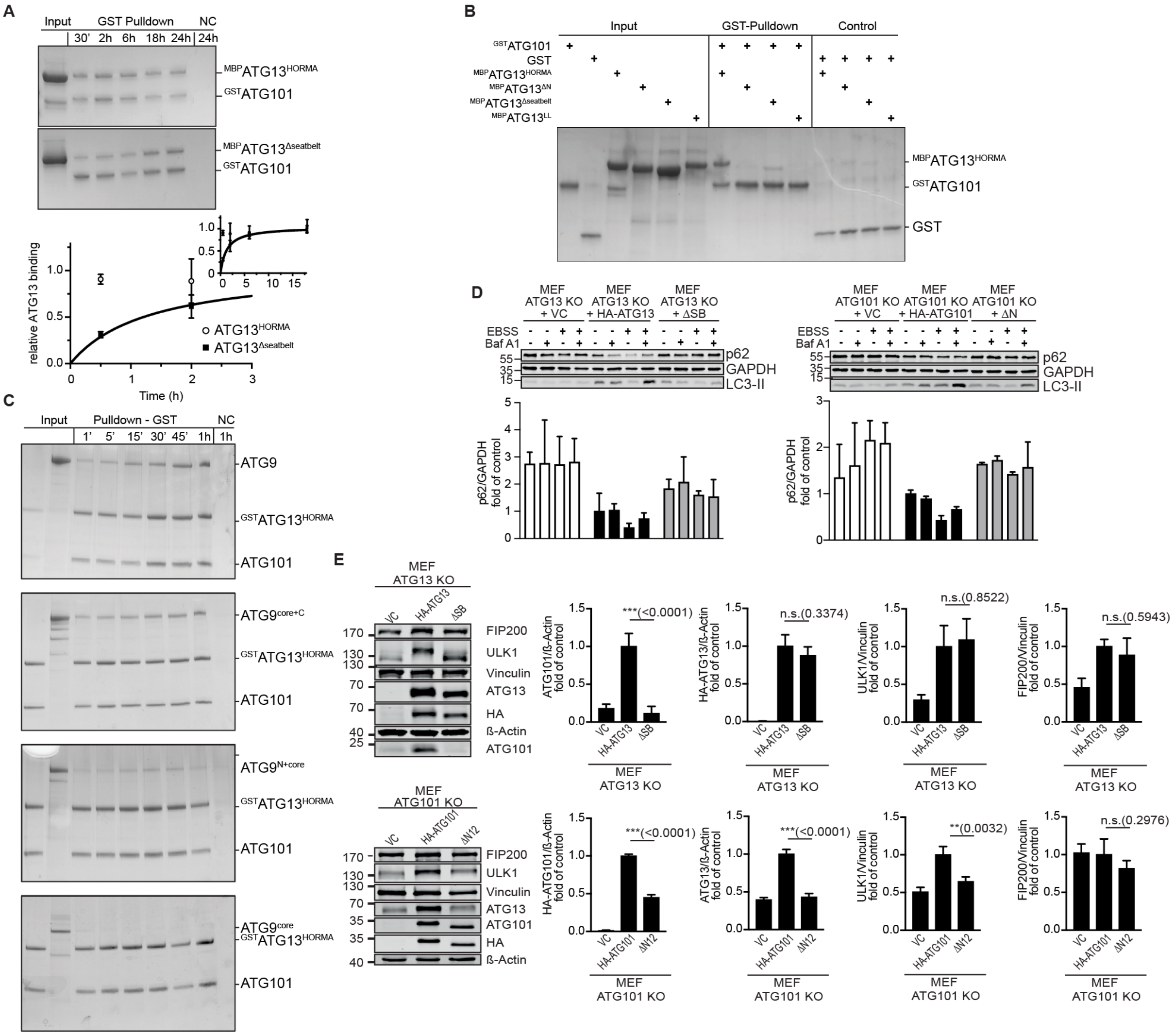


**Figure S3.** (**A**) The ATG13$^{\Delta seatbelt}$ mutant can interact with ATG101, but much slower than ATG13$^{WT}$. Pull-down experiment using $^{GST}$ATG101 as bait. Proteins were incubated at 20 °C for the time indicated. The assay was performed with 3 μM $^{MBP}$ATG13 and 1 μM of bait. The experiment was repeated at least three times with basically identical results. Quantification of the relative ATG13 binding is shown in the lower panel. (**B**) Mutating the mobile elements in ATG13 affects instantaneous ATG101 interaction. Pull-down experiment using $^{GST}$ATG101 as bait. The assay was performed with 3 μM $^{MBP}$ATG13 mutants and 1 μM of bait. Proteins were incubated at 4 °C for 1 h. (**C**) ATG9 truncations interact with the ATG13$^{HORMA}$-101 complex with different kinetics. Pull-down experiment using $^{GST}$ATG13$^{HORMA}$-101 as bait. Proteins were incubated at 4 °C for the time indicated. The assay was performed with 0.1 μM ATG13$^{HORMA}$-101 and 0.2 μM ATG9 truncations. (**D**) Mutants that modulate conformer state of ATG13 and ATG101 decrease basal and starvation-induced p62 turnover in MEFs. KO MEFs retrovirally transfected with empty vector (VC) or cDNA encoding the indicated ATG13 (left) or ATG101 (right) variant were incubated for 2 h in growth medium or starvation medium (EBSS) with or without 40 nM Bafilomycin A$_1$ (Baf A$_1$). ΔSB = Δseatbelt (Ext. data Fig. 1a). Western blots are the same as in Fig 2d,e. Results are mean + SD. Statistical analysis was performed using two-way ANOVA (corrected by Tukey‘s multiple comparisons test). (**E**) Mutating the mobile elements of the HORMA domain in ATG13 and ATG101 affects the stability of the ULK1 complex. Cleared cellular lysates of *Atg13* KO or *Atg101* KO MEFs retrovirally transfected with empty vector (VC) or cDNA encoding the indicated ATG13 or ATG101 variants were analyzed by immunoblotting for ATG101, ATG13, ULK1, FIP200, HA, ß-Actin and Vinculin. Fold changes were calculated by dividing each normalized ratio (protein to loading control) by the average of the ratios of the control lane (overexpressed HA-ATG13 or HA-ATG101, respectively). Results are mean + SD. Statistical analysis was performed using one-way ANOVA (corrected by Dunnett‘s multiple comparisons test). P values < 0.05 were considered statistically significant. Statistically significant/non-significant differences are only indicated for the wild-type vs. the variant protein. n.s. non-significant, *P < 0.05, **P < 0.01, ***P < 0.001, ****P < 0.0001.

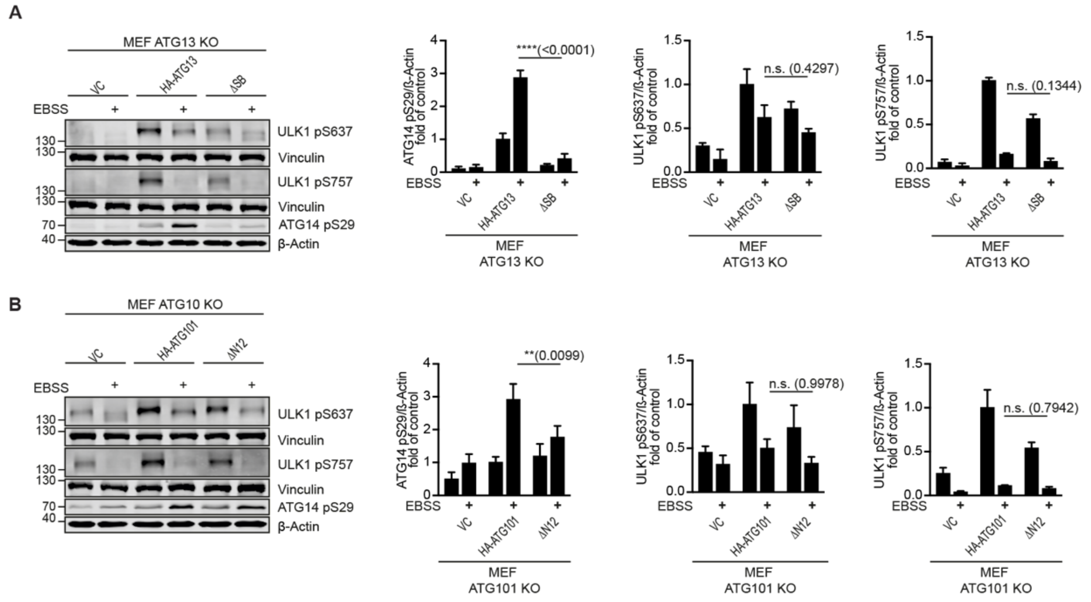


**Figure S4.** (**A**,**B**) Mutating the mobile elements of the HORMA domain in ATG13 (A) and ATG101 (B) affects the phosphorylation status of ULK1 and its substrate ATG14. *Atg13* KO or *Atg101* KO MEFs retrovirally transfected with empty vector (VC) or cDNA encoding the indicated ATG13 or ATG101 variants were incubated for 2 h in growth medium or starvation medium. Cleared cellular lysates were analyzed by immunoblotting for ULK1 phospho-S757 and S637 (corresponds to human S638), ATG14 phospho-S29, ß-Actin and Vinculin. Fold changes were calculated by dividing each normalized ratio (protein to loading control) by the average of the ratios of the control lane (overexpressed HA-ATG13 or HA-ATG101). Results are mean + SD. Statistical analysis was performed using two-way ANOVA (corrected by Tukey's multiple comparisons test; every mean was compared regardless to the treatment or the cell line). P values < 0.05 were considered statistically significant. Statistically significant/non-significant differences are only indicated for the wild-type vs. the variant protein in A-D. n.s. non-significant, *P < 0.05, **P < 0.01, ***P < 0.001, ****P < 0.0001.

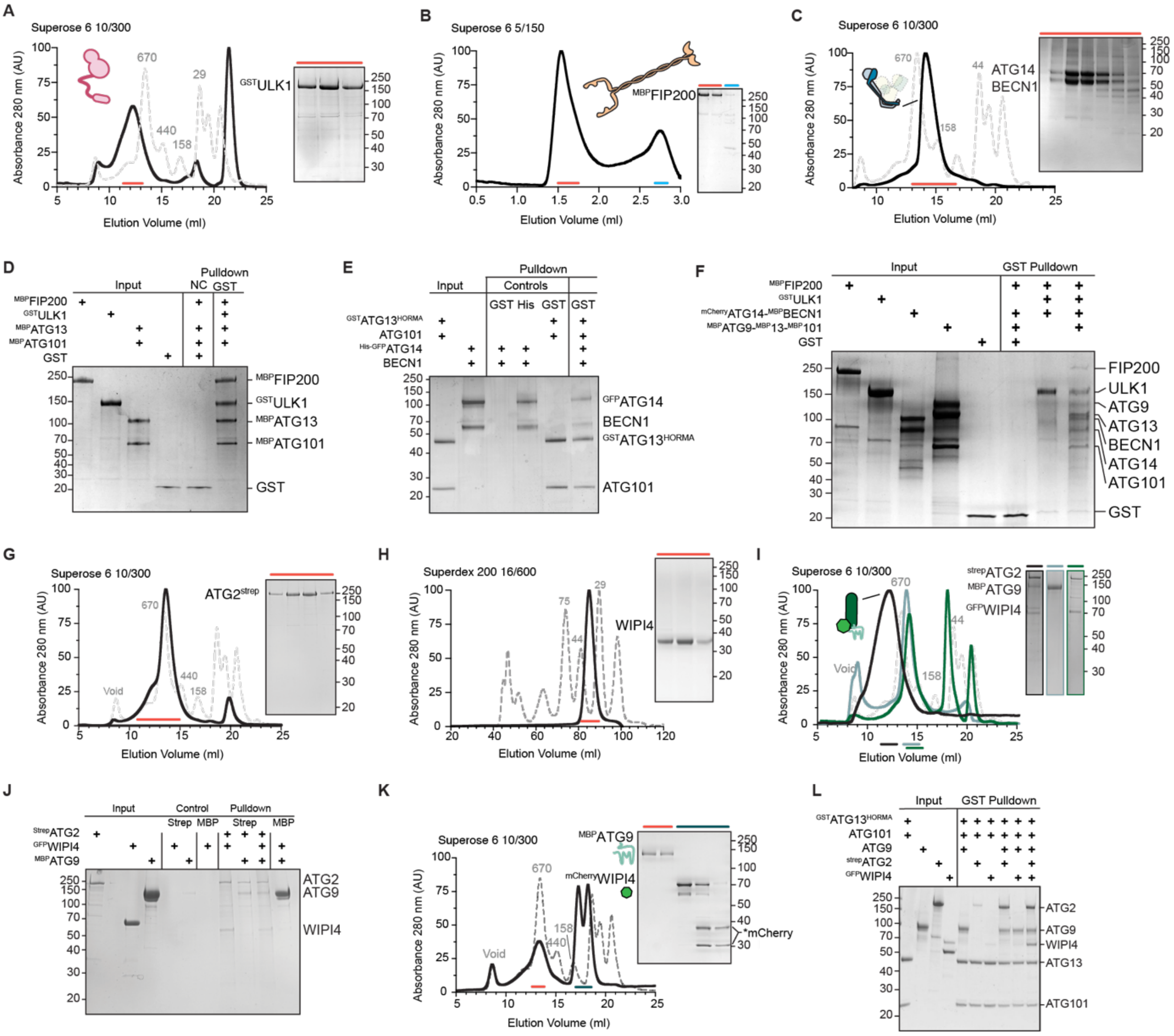


**Figure S5.** (**A**) SEC profiles of $^{\mathrm{GST}}$ULK1 (black trace), and marker proteins (gray dash-line trace) run on a Superose 6 10/300 SEC column and analyzed by SDS–PAGE. (**B**) Analytical SEC profiles of $^{\mathrm{MBP}}$FIP200 (black trace), and marker proteins (gray dash-line trace) run on a Superose 6 15/1500 SEC column and analyzed by SDS–PAGE. (**C**) SEC profiles of ATG14-BECN1 complex (black trace), and marker proteins (gray dash-line trace) run on a Superose 6 10/300 SEC column and analyzed by SDS–PAGE. (**D**) *In vitro* GST pull-down assay showing ULK1 complex consisting of 4 proteins: $^{\mathrm{MBP}}$FIP200- $^{\mathrm{GST}}$ULK1-$^{\mathrm{MBP}}$ATG13-$^{\mathrm{MBP}}$ATG101. The assay was performed with 1 μM of all proteins. (**E**) ATG14-BECN1 complex interacts with the ATG13$^{\mathrm{HORMA}}$-ATG101 complex. Pull-down experiment using $^{\mathrm{GST\text{-}}}$ATG13$^{\mathrm{HORMA}}$-ATG101 as bait. Proteins were incubated at 4 °C for 1 h. The assay was performed with 1 μM ATG13$^{\mathrm{HORMA}}$-ATG101 and 3 μM ATG14-BECN1. (**F**) Without ATG9-13-101 the 7-subunit initiation super-complex does not form. Pull-down using $^{\mathrm{GST}}$ULK1 was performed with 1 μM of all protein. Pull-down was performed using 1 μM bait and 3 μM prey, with the proteins were incubated at room temperature 1 h. (**G**) SEC profiles of ATG2$^{\mathrm{Strep}}$ (black trace), and marker proteins (gray dash-line trace) separated using a Superose 6 10/300 SEC column and analyzed by SDS–PAGE. (**H**) SEC profiles of WIPI4 (black trace), and marker proteins (gray dash-line trace) separated using on a Superdex 200 16/600 SEC column and analyzed by SDS–PAGE. (**I**) SEC profiles of $^{\mathrm{MBP}}$ATG9-ATG2$^{\mathrm{Strep}}$-$^{\mathrm{GFP}}$WIPI4 complex (black trace), ATG2$^{\mathrm{Strep}}$-$^{\mathrm{GFP}}$WIPI4 complex (green trace), $^{\mathrm{MBP}}$ATG9 (blue trace), and marker proteins (gray dash-line trace) run on a Superose 6 10/300 SEC column and analyzed by SDS–PAGE. Analysis of corresponding peak fractions demonstrates co-elution of the subunits. (**J**) SEC profiles of a mixture of ATG9 and WIPI4 (black trace), and marker proteins (gray dash-line trace) run on a Superose 6 10/300 SEC column and analyzed by SDS–PAGE. The two proteins did not co-elute, indicating no interaction. (**K**) *In vitro* GST and Strep pull-down assays showing ATG2-WIPI4-ATG9 complex. ATG2 interacted with both WIPI4 and ATG9, however ATG9 did not interact with WIPI4. Pull-down was performed using 1 μM bait and 3 μM prey, with the proteins were incubated at room temperature 1 h. (**L**) ATG9-13-101 complex interacts with the ATG2-WIPI4 complex. Pull-down experiment using $^{\mathrm{GST}}$ATG13$^{\mathrm{HORMA}}$-ATG101 as bait. Proteins were incubated at 4 °C for 1 h. The assay was performed with 1 μM of bait and 3 μM of prey.

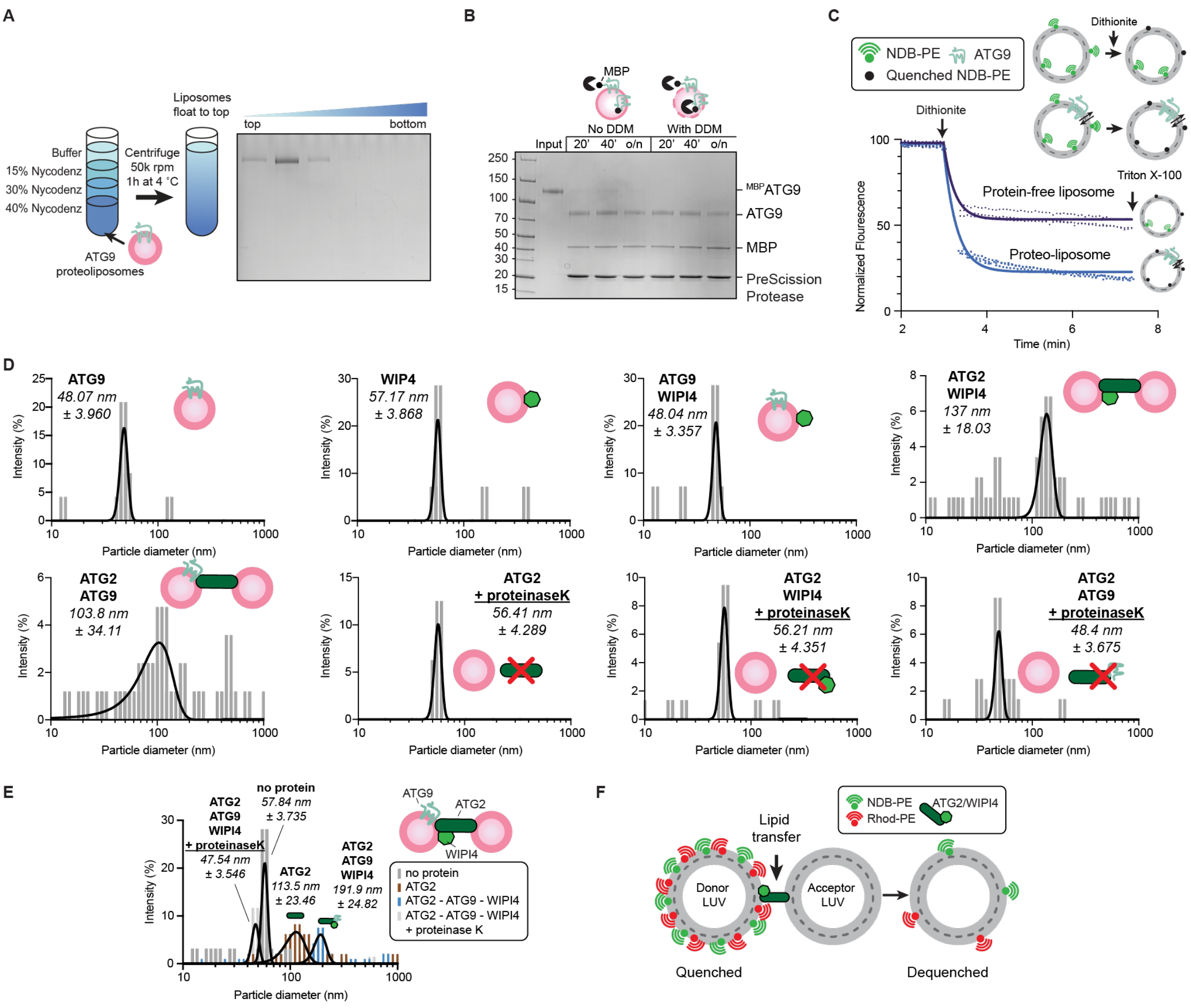


**Figure S6.** (**A**) LUVs reconstituted with ATG9 were subjected to Nycodenz step gradient. After ultracentrifugation, the gradient was fractionated and analyzed by SDS. ATG9 proteoliposomes were only found in the top fractions indicating successful reconstitution. (**B**) Protease protection assay of ATG9 proteoliposome showing almost complete cleavage of the N-terminal MBP tag of the protein compared to the control in the presence of DDM, indicating that the N-terminal cleavage site of ATG9 was oriented outwards after reconstitution. (**C**) Compared to protein-free control, the ATG9-proteoliposome showed significantly higher quenching level of NBD signals, indicating scramblase activity of ATG9. Schematic of a scramblase assay described in methods section. **D**) DLS profiles showing size distribution of LUVs in the presence or absence of indicated proteins. All soluble proteins are added to a final concentration of 200 nM. ATG9 PL is added at a final lipid concentration of 50 µM at a protein to lipid ratio of 1:2000. (**E**) DLS profiles showing size distribution of LUVs in the presence of ATG2 (brown); ATG2, ATG9, WIPI4 (blue); no protein (dark gray); or ATG2, ATG9, WIPI4. After incubation with Proteinase K (dark gray). A shift to larger size in the presence of indicated protein indicates tethering activities. A shift to smaller size in proteinase K treatment indicates tethering activities are protein-induced and not caused by vesicle fusion. (**F**) Schematic of the lipid transfer assay described in the methods section.
